# Microbial activity transitions underlie community stability following persistent warming

**DOI:** 10.64898/2026.11.26.768007

**Authors:** Xipeng Liu, A. Fina Bintarti, Margaux Buron, Milena Gonzalo, Ashley Shade

**Affiliations:** Université Lyon 1, CNRS, INRAE, LEM, UMR 5557, Villeurbanne, France

## Abstract

Microbial responses to environmental disturbances are often assessed through community composition or bulk biomass, but whether physiological state changes provide complementary and indicative information about community stability remains unclear. By tracking cell abundance, redox activity, 16S rRNA:rRNA gene ratios, soil respiration, and stability across six soil communities spanning three ecosystem types and two seasonal histories under persistent warming, we found that cell abundance and physiological activity exhibited asynchronous dynamics across biological levels. Early physiological activity transitions, compared to abundance changes, exhibit greater interpretability based on their pre-perturbation state. Crucially, early activity transitions predicted subsequent stability at population and community levels, whereas abundance-based metrics lacked comparable predictive power. Our findings establish physiological activity transitions as an early, essential dimension connecting environmental history to microbial community trajectories, offering a predictive framework for forecasting microbiome stability under global change.

## 1 Introduction

Soil microorganisms drive a large fraction of terrestrial organic matter decomposition and mediate the transformation and retention of essential nutrients, thereby linking environmental change to ecosystem functioning and climate–carbon feedbacks^1–3^. Microbial responses to environmental change span multiple dimensions, including population size, taxonomic identity and physiological activity state, each capturing a distinct facet of community dynamics^4–6^ in which cellular and population responses nested in the community level. Yet these dimensions need not change in concert, and the coordination of their trajectories across different biological levels during environmental change remains poorly understood. Consequently, measurements restricted to a single dimension may overlook crucial components of microbial responses and lead to incomplete interpretations. Resolving these complementary dimensions is therefore essential for understanding responses of microbial communities and ecosystem stability to a changing climate.

Warming is a key environmental factor structuring microbial communities in the context of climate change. Temperature directly alters biochemical reaction rates, growth and respiration, while sustained warming can impose selection on microorganisms with different thermal niches and resource-use strategies^7–10^. Responses to temperature can therefore emerge at multiple biological levels, from rapid physiological adjustment within cells to changes in the abundance and thereby representation of populations within communities. Experimental studies of individual bacterial species, for example, have shown that prolonged exposure to elevated temperature can alter respiratory temperature sensitivity and growth optima, indicating that microbial populations can adjust their thermal traits under persistent warming^11,12^. At the community level, warming can restructure microbial composition and selectively favor populations with higher thermal tolerance, growth potential or stress resistance^13–15^. Importantly, community-level responses need not result from uniformly accelerated growth of individual populations: long-term warming has been shown to increase community growth primarily by increasing the number of growing bacterial taxa rather than their taxon-specific growth rates^16^. Conversely, persistent warming can also reduce microbial biomass as substrate depletion constrains microbial activity and growth^17^. Together, these findings indicate that warming can reshape microbial communities through changes in physiological state, and that population abundance and community composition that may follow distinct trajectories (Fig. 1a). A subsequent but unresolved question is whether early physiological state transitions contain indicative information about longer-term community stability beyond that provided by abundance and composition. Yet most studies have emphasized relative community composition and bulk biomass, leaving the dynamics of absolute microbial abundance and physiological activity across cellular, population and community levels comparatively unresolved.

**Fig. 1.**
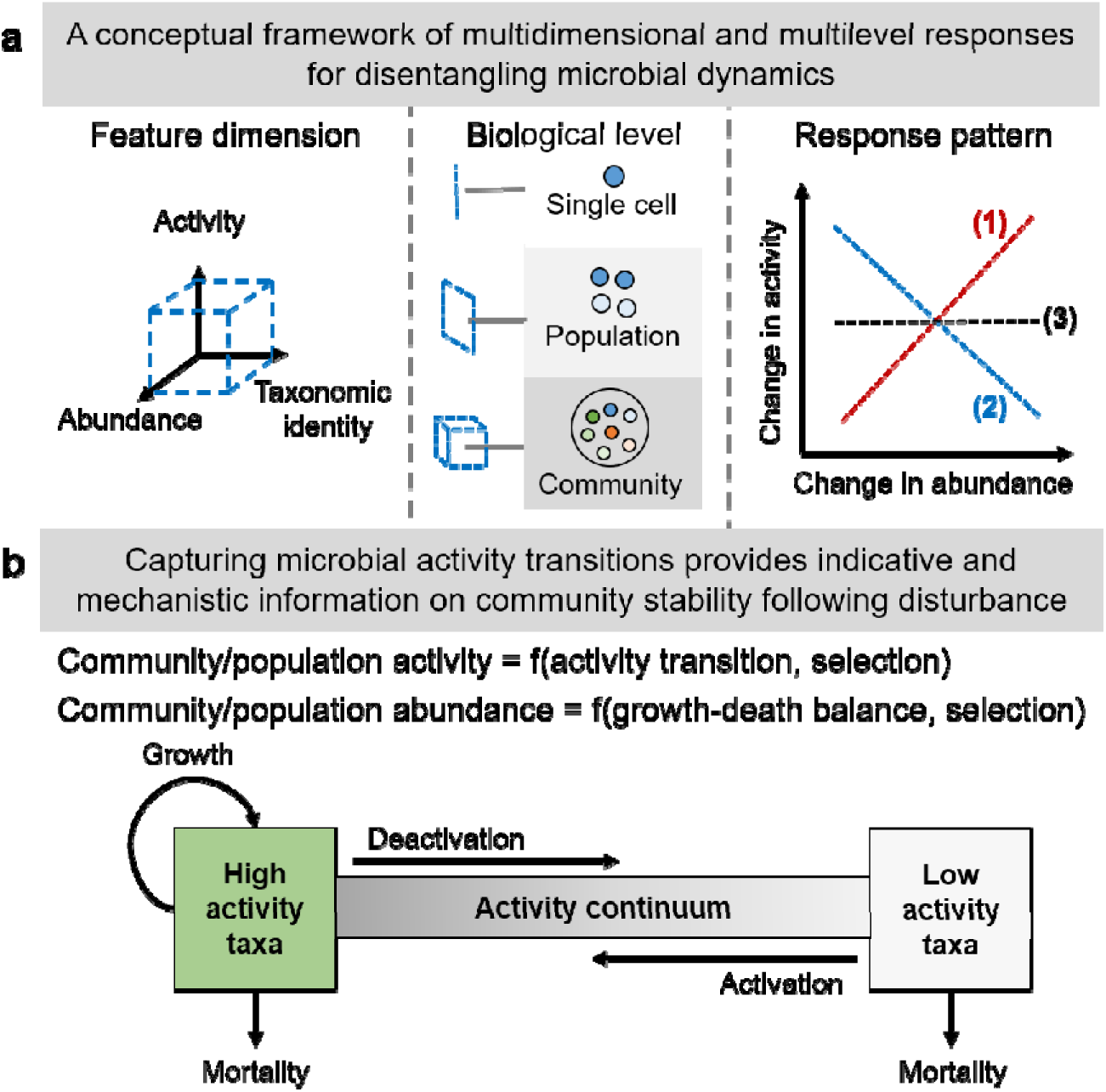
Conceptual framework integrating microbial abundance, activity, and composition for understanding community dynamics following disturbance. **(a)** Microbial community responses to environmental disturbance can be characterized by three complementary dimensions: physiological activity, demographic abundance, and taxonomic composition. Activity and abundance can exhibit three broad patterns of change at both population and community levels: they can change in (1) the same direction, (2) change in opposite directions, or (3) change in one dimension while remaining relatively stable in the other. The latter two patterns indicate the asynchronous dynamics between microbial activity and abundance. Community composition, in turn, captures changes in the identity and representation of microbial populations. **(b)** Conceptual linkage between microbial physiological activity transitions and demographic and compositional dynamics. Along the activity continuum, microbial activation and deactivation are shaped by deterministic selection imposed by biotic and abiotic factors, with stochastic processes driving variation in microbial physiological states. Population abundance changes consequently reflect the balance between growth and death, while changes in community membership reflect shifts in population representation. Integrating these processes links physiological responses to demographic and compositional reorganization and provides a basis for understanding and predicting community stability.

The microbial dormancy framework provides a conceptual starting point for linking physiological state to community dynamics^18,19^, traditionally describing populations as comprising active and dormant fractions that can reversibly transition between states in response to environmental conditions. However, microbial activity is increasingly recognized as a continuum, ranging from highly active and dividing cells through metabolically active but non-dividing states to deeply dormant phenotypes^20^. We view this continuum as a fundamental property of individual cells that can be scaled across levels of biological organization: at the population level, it is represented by the distribution of physiological states across constituent cells, and at the community level, by the aggregate distribution of these states across populations (Fig. 1a). Environmental perturbations can therefore shift physiological activity along this continuum toward either higher or lower activity without requiring transitions between discrete active and dormant categories (Fig. 1b). Moreover, activity transitions provide a physiological dimension for connecting the enrichment, depletion, persistence, or replacement of cells and populations, while changes in abundance and membership describe the demographic and compositional consequences of these shifts. From this perspective, activity transitions are essential for a unified conceptual framework to describe how microbial communities reassemble over time (Fig. 1b). This framework therefore motivates examining the response patterns of microbial activity to environmental perturbation and how early activity states and transitions contribute to the subsequent stability of microbial communities (Fig. 1).

Here, we tested the response pattern between activity and abundance and examined the information provided by early microbial physiological activity states for community stability following persistent warming. We tracked cell abundance, redox activity, 16S rRNA:rRNA gene ratios, community composition, and CO□emission across six soil communities differing in ecosystem type and seasonal history. We hypothesized that microbial abundance, physiological activity, and composition would respond along distinct trajectories, and that early activity transitions during the warming would be more predictive of subsequent population and community stability than abundance-based changes alone.

## 2 Methods

### Soil sampling and persistent warming response experiment

Soils used in this study were collected from three ecosystem types (Agriculture, Grassland, and Forest) in France and from winter (January and February, 2025) and summer (June and August, 2025) seasons, respectively. This results in six soils in total, including three different soils from two temperature levels (Cold and Warm). Soil temperature ranges from 4-7 °C in winter, and from 19.2-20.5 °C in summer, and the basic properties of soils were distinct across sites (Supplementary Table S1). For the soil sampling, we used clean metal shovels or soil drills to collect soil from a depth of 0-20 cm, removing plant and animal residue, and recording the soil temperature at the time of collection. After collection, the soil was quickly transported back to the laboratory, passed through a sieve with a 4-mm aperture, and stored in a constant-temperature incubator (New Brunswick Innova 42R, Eppendorf) at the origin soil temperature for 3-7 days.

The microcosms were made by adding 1200 g of sieved soil into an autoclaved glass jars with lids that were not fully closed to prevent anoxia. The jars were transitioned from the soil origin temperatures into an incubator set to 25 °C, arranged randomly, and allowed to stabilize over 27 days. This design allowed us to examine microbial responses under a consistent experimental temperature while retaining differences in the initial environmental history and warming intensity experienced by communities from different seasons and ecosystems. The total mass of each microcosm was recorded at the beginning of experiment and after each sampling to track soil moisture changes. Soil moisture was kept constant during the incubation. Four biological replicates were included for each soil, resulting in a total of 24 microcosms.

To monitor the changes in microbial communities, we conducted non-destructive sampling from each microcosm at seven time points: days 0, 3, 6, 9, 15, 21, and 27 after incubation. Day 0 represented the pre-warming baseline, while Day 3 captured the early transition to persistent warming conditions. Briefly, 30.0 g of soil was collected at each time point using a small metal core (1.5 cm in diameter and 20 cm long) by coring three to four times within each microcosm. After sampling, the sampled holes were gently filled with the surrounding soil and the microcosms were immediately placed back into the incubator. The masses of microcosms were recorded before and after soil sampling to monitor potential water evaporation. The soil moisture content was maintained at its Day 0 initial state by adding sterile water.

### Soil microbial abundance and redox activity

The cell abundance and redox activity of soil microbial communities were measured using our previously developed method^21^, which consisted of cell extraction from the soil, staining, detection, and automated image analysis (see Supplementary Methods). The abundance of total cells (intact cells) and active cells with detectable redox activity in each community is expressed as cell count per gram of dry soil. Fluorescence intensity reflects redox activity and is used to infer the redox activity of the community.

### Soil CO_2_ emission capacity

Bulk soil CO_2_ emission was measured using a gas chromatograph (Micro GC R3000, SRA Instrument, Marcy L’Etoile, France) as previously described^22^. In brief, 15.0 g of freshly collected soil was placed in a new glass plasma flask that was hermetically sealed with a rubber stopper. After a thirty-minute acclimation, automatic measurements were taken every 30 min for 7 h at 25 °C. The data from forest soils (Days 9 and 27) and grassland soils (Days 9 and 27) were not recorded and therefore excluded from the following analyses due to unexpected reasons during measurement.

Soil CO□emission dynamics were modeled using a nonlinear Bayesian model implemented in the “brms” package^23^ (v. 2.23.0) in R. For each experiment, CO□accumulation over incubation time was described using a modified asymptotic respiration model:

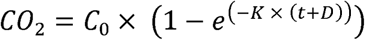

where *C_0_* represents the limiting substrate at the beginning of the experiment, *K* represents the respiration rate constant, *t* is the calibrated incubation time, and *D* represents a time-shift parameter accounting for variation in the initial phase of CO□accumulation. CO□observations were modeled using a Gaussian likelihood^21^. Model convergence and fit were assessed using posterior summaries and graphical posterior predictive checks. To quantify the initial respiration rate, posterior draws of C, K, and D were extracted with “tidybayes” package^24^ (v. 3.0.7) and combined with the posterior distributions of the experimental-unit-specific random effects. The initial capacity of soil CO_2_ emission (i.e., CO_2_ emission capacity) was calculated and expressed as the amount of carbon dioxide (ppm) produced per gram of soil dry weight per hour.

### Microbiota profiling

A detailed description of the following standard protocols is provided in Supplementary Methods. Total nucleic acids (DNA and RNA) were co-extracted from 0.5 g of soil following a modified protocol based on Griffiths et al. (2000)^25^, with the RNA component purified using DNAse treatment and then synthesized into cDNA. Illumina sequencing of 16S rRNA and 16S rRNA gene (also known as rDNA) was performed targeting the V3-V4 region using standard library preparation protocols. Amplicon sequencing data targeting were processed using QIIME 2 (v. 2026.4)^26^ according to standard pipelines. OTUs were defined at 99% sequence identity to account for possible amplification errors introduced by the reverse transcription step of the rRNA to cDNA that could ultimately create artificial non-coherence in the rRNA and rDNA-defined taxa.

### Population abundances

To account for variation in the number of 16S rRNA gene copies among bacterial taxa, OTU abundances of rRNA and rDNA tables were corrected using predicted 16S rRNA gene copy numbers. Representative OTU sequences were exported from QIIME 2 and used as input for “RasperGade16S”^27^, which predicts taxon-specific 16S rRNA gene copy numbers directly from 16S rRNA gene sequences. Predicted copy numbers were obtained for all representative OTUs and retained for subsequent correction of DNA-based OTU abundances. This correction was applied to reduce biases in estimates of bacterial abundance arising from interspecific variation in 16S rRNA operon copy number. The copy-number corrected OTU abundance matrix was relativized to obtain relative tables from rDNA and rRNA. The relative abundances of rDNA and rRNA were considered as proxy for relative genomic abundance and ribosomal abundance, respectively. We then assessed the response patterns of these two metrics as described below. Moreover, the absolute cell abundance matrix for each sample was obtained by multiplying by the image-quantified total number of cells detected per gram of soil.

### Population-level relative ribosomal activity and activity transitions

#### Calculation

Because 16S rRNA abundance does not directly measure metabolic flux or realized metabolic rates of a single population^28,29^, we used the ratio of 16S rRNA to 16S rRNA gene abundance (16S ratio) as a proxy for the relative activity state, representing the relative ribosome investment of the bacterial population. For each OTU in each sample, the 16S ratio was calculated as the abundance of the rRNA-derived amplicon divided by the corresponding rDNA-derived gene abundance^30^.

### Population-level activity classifications

Based on this ratio and the detection status of each OTU in the rDNA and rRNA datasets, out-sample combinations were categorized into one of four categories of metabolic potential, representing an activity gradient. OTUs detected in the rDNA fraction were classified as high metabolic potential when the 16S ratio was ≥1, moderate metabolic potential when the ratio was ≥0.25 and <1, and low metabolic potential when the ratio was <0.25. OTUs with a detectable rRNA signal but no detectable rDNA signal were classified as “phantom” OTUs^31^, reflecting the absence of a corresponding detectable genomic signal. Notably, the 16S ratio was applied a continuous measure of relative ribosomal activity without applying any ratio threshold for exclusion, and the categories were used only to facilitate comparison and visualization of population-level activity patterns. Note that the thresholds of 1 and 0.25 were defined as operational categorical thresholds based on the observed distributions of 16S ratios in our dataset, rather than as universal physiological or ecological boundaries. Accordingly, these categories should be interpreted as relative states of transcriptional investment normalized to genomic abundance within the present dataset, rather than as discrete levels of realized metabolic activity.

### Activity transitions

The change in 16S ratio of each OTU across time was interpreted as the transition of population-level ribosomal activity. We used this metric to capture temporal changes in the relative ribosomal activity of individual populations while accounting for their corresponding 16S rRNA gene abundance. Temporal changes in the ratio were therefore used to distinguish shifts in population activity from changes in population size, with positive and negative values indicating increases and decreases in relative ribosomal activity, respectively. This approach enabled us to quantify the temporal dynamics of microbial activity at the population level following warming.

### Quantifying stability

The two sets of stability quantifications included integrated and component-level measures^32^. Temporal stability represents the integrated measure and was quantified as the inverse of the coefficient of variation (CV), which indicates the invariability of the parameter of interest (e.g., community and population cell abundance, redox activity, rRNA and rDNA relative abundances, relative ribosomal activity) over time, relative to the relative long-term average level:

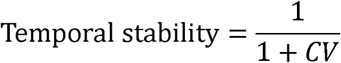

Temporal stability was calculated for the entire study period (Day 0 to Day 27) and for the later stage (Day 3 to Day 27) that excluded the early period (Day 0 to Day 3).

Resistance was quantified as the degree to which a response variable remained close to its non-disturbed reference state (i.e., Day 0). Resistance was calculated as the complement of the relative displacement from the Day 0:

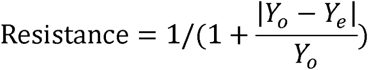

where *Y_O_* represents the response variable at Day 0 and *Y_e_* represents the corresponding value for an individual replicate at the specified post-disturbance time point. Thus, a resistance value of 1 indicates no deviation from the non-disturbed state, whereas decreasing values indicate increasing displacement from the Day 0 reference state. Similarly, a stability value of 1 indicates no temporal variation across the observation period, whereas lower values indicate greater temporal variability relative to the temporal mean and therefore lower stability. Resistance was evaluated at Day 3 and Day 27 to characterize relatively short-term and long-term responses to persistent disturbance, respectively, and the resistance and temporal stability at population and community levels were accordingly calculated.

Community compositional resistance and stability were quantified based on community composition similarity across time. Weighted UniFrac distance was calculated between consecutive sampling points for each replicate to quantify compositional displacement over time. Resistance was evaluated at Day 3 and Day 27, respectively. The mean temporal distance (D^_^) across all consecutive intervals was then used as a measure of temporal compositional variability. To express stability on a positive scale, compositional stability was calculated as the inverse of temporal displacement:

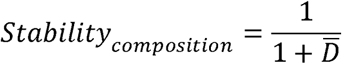

### Statistical analysis

All statistical analyses were conducted in R version 4.5.2. Differences in resistance, temporal stability, and community succession, among sampling seasons and site types were assessed using linear models or linear mixed-effects models, as appropriate for the data structure, as the response and season, site type, and their interaction as fixed effects. Model effects were evaluated using analysis of variance (ANOVA), and model *R*^2^ was calculated to quantify the variance explained. Where significant effects were detected, pairwise differences among site types within each season were evaluated using estimated marginal means with Tukey adjustment for multiple comparisons. Differences in community composition were tested using PERMANOVA (adonis2) based on weighted UniFrac distances calculated from the OTU absolute abundance table, with season, site type, sampling day, and their interactions as explanatory variables; 999 permutations were performed within microcosm trajectories to account for the repeated-measures structure. To capture short-term ecological variability and compositional turnover, we computed rolling rates of change for each response variable within each experimental unit. For population abundances and redox activity, we calculated the relative mean squared successive difference (MSSD) using R package “slider” (v. 0.3.3)^33^, i.e. the windowed mean of squared rates of change normalized by the squared rolling mean, yielding a dimensionless index of temporal instability.

Changes in community- and OTU-level metrics were quantified as the log_2_ fold change between consecutive sampling time points. The relationships between changes in demographic and physiological metrics across population and community levels were assessed with linear models or linear mixed-effects models, with season and site type as categorical predictors, and all their two- and three-way interactions. Significant differences in slopes among season–site combinations were further tested using estimated marginal trends with R package “emmeans”^34^ (v. 2.0.1), with Tukey adjustment for multiple comparisons. OTUs with a 16S ratio or relative abundances of zero are not included in this analysis because the true log fold change cannot be precisely calculated. Furthermore, we quantified temporal trajectory asynchrony between rRNA and rDNA trajectories at the OTU level using dynamic time warping (DTW), which identifies the dissimilarity between two standardized trajectories. DTW distance was calculated between the Z-score-standardized trajectories using R package “dtw” (v. 1.23)^35^, with a Sakoe–Chiba window size of zero to constrain alignment to corresponding time points. This index was normalized to a common range between 0 (full synchronous changes) and 1 (full asynchronous changes) using the global minimum and maximum values for each metric, while preserving the relative ranking of OTUs.

To assess early transitions of population states, changes in OTU absolute abundance and 16S ratio were evaluated between Days 0 and 3. OTUs detected in at least two biological replicates at only one of the two time points were classified as activated or deactivated based on 16S ratio, or increased or decreased based on absolute abundance, according to the direction of change. For OTUs detected at both time points, the significance of changes was assessed using Welch’s %-tests on log2-transformed values. The count, proportion, initial cell abundance, and taxonomic identity of OTUs that significantly changed in activity or abundance were identified.

To examine whether initial states and early responses predicted (subsequent) stability at community and population levels, we used partial least squares structural equation modeling (PLS-SEM) to examine the direct and indirect relationships among initial community states, transitional dynamics, and community/population stability at cell abundance, physiological activity, and composition dimensions. For each response domain, we specified a separate PLS-SEM consisting of a measurement model and a structural model and all models were estimated with the “seminr” package^36^ (v. 2.5.0) with default settings for the PLS algorithm. Multiple observed variables representing the same conceptual domain were combined into composite constructs, with indicator assignments for initial states, transitional dynamics, and stability specified separately for each response domain and analytical level. The complete model specifications, including indicator assignments and parameter settings of each model, are provided in the Supplementary Table 2. To obtain standard errors, confidence intervals, and significance tests for path coefficients without relying on distributional assumptions, we performed bootstrapping with 10000 resamples. Bias-corrected and accelerated (BCa) 95% confidence intervals were computed, and paths were considered significant if their confidence intervals excluded zero (α = 0.05).

## Data availability

The sequence data generated in this study have been deposited in the NCBI Sequence Read Archive under BioProject number PRJNA1554522. Microscope images for total and active cells are available on FigShare (10.6084/m9.figshare.34768650).

## Code availability

All code for the bioinformatic sequence processing in QIIME 2 and statistical analysis in R is available on GitHub (https://github.com/ShadeLab/Stability_MicroRescue_Liu_2026). Code for automated image processing is available (https://github.com/DatInsights-com/process_quantom).

## 3 Results

### Warming produced distinct responses in microbial abundance, activity, and composition

Microbial abundance, activity, and community composition showed distinct responses to warming across ecosystem types and seasons (Fig. 2, Supplementary Fig. 1, and Supplementary Text 1), indicating that these dimensions captured complementary aspects of microbial community dynamics. Active and total cell abundances differed substantially among ecosystem types, with forest soils generally supporting larger bacterial populations (10^9^–10^10^ cells g^-1^ dry soil) than grassland and agricultural soils (10^8^–10^9^ active cells g^-1^ dry soil). Active-cell-abundance resistance was strongly context dependent during the early response: at Day 3, season, site type, and their interaction were all significant (*P* = 0.014, 0.0060, and 0.0037, respectively; adjusted *R*² = 0.58). Under cold-season conditions, agriculture and grassland soils showed higher resistance than forest soils, whereas soil-type differences were absent under warm-season conditions. By Day 27, season and site type remained significant (*P* = 0.034 and 0.00073; adjusted *R*² = 0.50), but their interaction was no longer significant, while temporal stability of active cell abundance was comparatively insensitive to season and site type (all *P* ≥ 0.05; Fig. 2a).

**Fig. 2.**
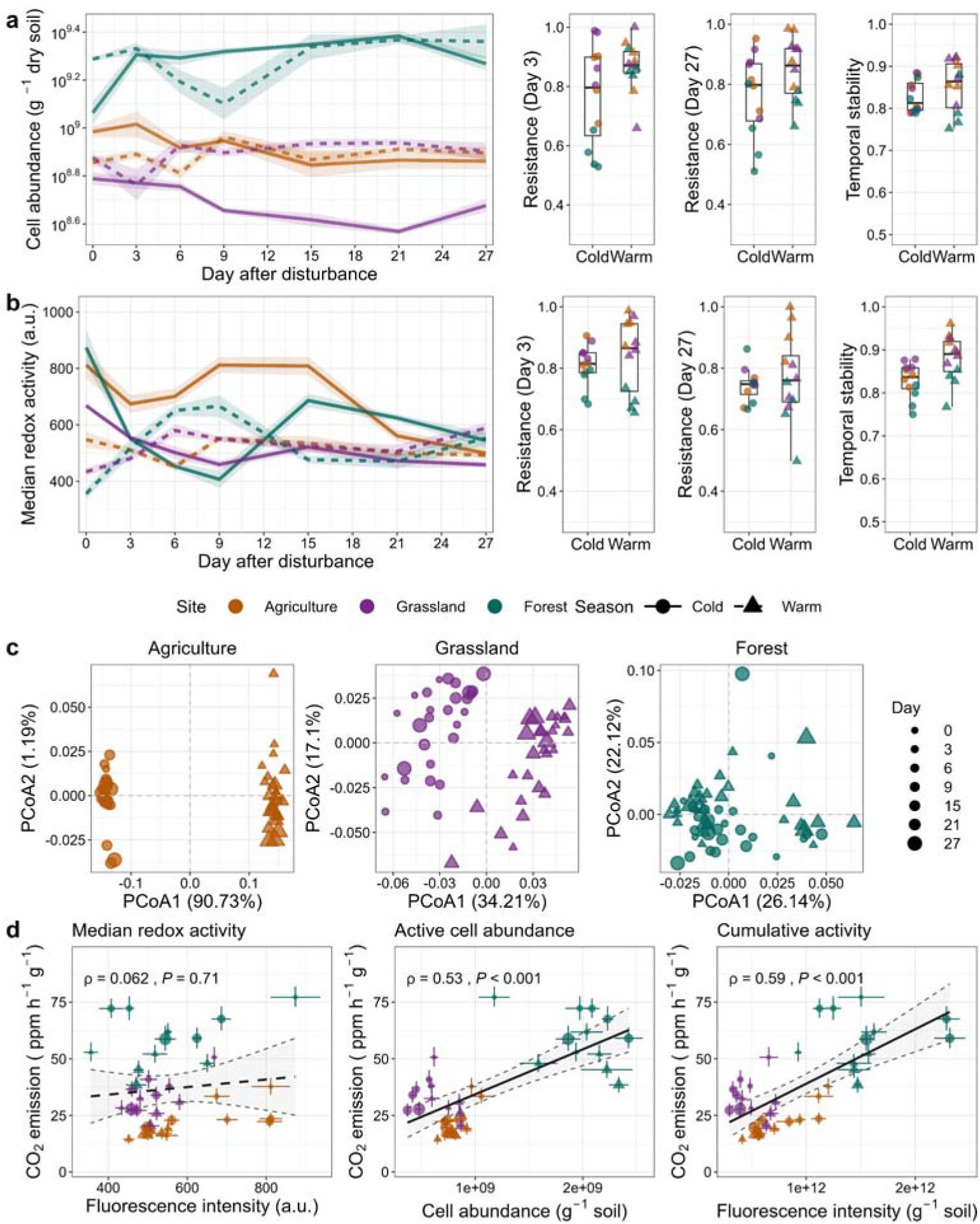
Microbial community dynamics under persistent warming. Dynamic, resistance, and temporal stability of active cell abundance (**a**) and community median redox activity (**b**) in soils. The shading around the line represents the standard error (n=4). (**c**) Beta diversity showing the succession of community composition across time in different sites. Principal Coordinate Analysis (PCoA) was based on weighted UniFrac distance. (**d**) Relationships between multiple dimensions of microbial community metrics and soil community functioning (CO□ emission rate) following disturbance. Spearman rank correlation was shown as regression lines and corresponding 95% confidence intervals. Vertical error bars of points indicate 95% credible intervals of CO□ emission derived from the posterior distribution using a Bayesian model, whereas horizontal error bars represent ±1 standard error of microbial community properties (n=4).

Community redox activity followed a contrasting trajectory. Initial median redox activity was substantially higher before warming in cold-season soils (654–985 a.u.) than in warm-season soils (304–617 a.u.), but subsequently converged toward a narrower range of activity levels (Fig. 2b). Unlike active-cell-abundance stability, temporal stability of community redox activity was strongly structured by environmental context, with significant effects of season and site type (*F* = 23.34 and 23.41, respectively; *P* = 0.0001 and < 0.0001; adjusted *R*² = 0.76). At Day 3, community activity resistance was primarily structured by site type (*F* = 32.95, *P* < 0.0001; adjusted *R*² = 0.77), with grassland soils showing higher resistance than forest soils under both seasonal histories. By Day 27, the response became strongly site dependent, as indicated by a significant season × site type interaction (*F* = 8.30, *P* = 0.0028; adjusted *R*² = 0.52): agricultural soils showed higher resistance than both forest and grassland soils under warm-season conditions, whereas no soil-type differences remained under cold-season conditions.

Community composition exhibited yet another response pattern. Weighted UniFrac composition was strongly differentiated by site type (PERMANOVA, *R*² = 0.70, *P* = 0.001) and seasonal history (*R*² = 0.073, *P* = 0.001), with a significant season × site type interaction (*R*² = 0.153, *P* = 0.001, Fig. 2c). Community composition also changed significantly through time, with significant day × season and day × site type interactions (*P* = 0.008 and 0.021, respectively), indicating that the trajectories of community restructuring differed among seasonal histories and soil types. The three-way season × site type × day interaction was not significant (*P* = 0.13), suggesting that these temporal effects were not strongly dependent on the joint combination of seasonal history and soil type (Fig. 2c).

Short-term temporal variability was highest immediately after warming and declined progressively over the 27-day period across microbial abundance, activity, and community composition (Supplementary Fig. 2). The four response variables differed markedly in how quickly they stabilized after warming. Both total and active-cell abundance showed a slower and more prolonged decline in relative rolling MSSD, remaining elevated for roughly the first 9–15 days before approaching zero. In contrast, community redox activity declined more rapidly and stabilized earlier, indicating that activity variability was resolved sooner than abundance variability. Community composition showed the least pronounced temporal signal overall, with only a gradual decline in rolling succession rate. These contrasts suggest that abundance and activity metrics capture different timescales of community response to warming.

Together, these results demonstrate that abundance, activity, and composition responded to warming through distinct combinations of initial state, resistance, temporal stability, and community restructuring. In particular, the contrasting stability patterns of active-cell abundance and community activity, together with the independent temporal restructuring of community composition, indicate that activity captures information that is not redundant with abundance or composition.

### Roles of demographic and physiological states in community functioning

Posterior predictive checks indicated that the fitted respiration model adequately reproduced the observed response distribution and average values (Supplementary Fig. 3). CO□ emissions declined rapidly during the initial phase following disturbance and subsequently stabilized, with clear differences among sites (Supplementary Fig. 4a). Community size (number of cells) and physiological activity contributed largely independently to variance in community respiration capacity: variance partitioning showed that active cell abundance uniquely explained 51% of the variation, whereas median redox activity uniquely explained 1%, with no detectable overlap between the two predictors (Supplementary Fig. 4b). Using total cell abundance yielded a similar pattern, indicating that community cell abundance and physiological activity represent complementary dimensions of microbial control over soil respiration, with cell abundance explaining the larger share of variance in soil CO□ emission. Consistent with this pattern, CO□ emission was positively associated with both active-cell abundance (ρ = 0.53, *P* < 0.001) and total cell abundance (ρ = 0.47, *P* = 0.003) and whereas instantaneous median redox activity was not associated with heterotrophic respiration (*P* = 0.71, respectively; Fig. 2d; Supplementary Fig. 4c,d). In contrast, cumulative bacterial activity showed the strongest positive association with CO□ emission (ρ = 0.59, *P* < 0.0001; Fig. 2d). Together, these results indicate that physiological state provides additional information for predicting community-level functioning beyond that captured by microbial community size alone, particularly when activity is integrated over time.

### Demographic and physiological responses are asynchronous

If physiological activity transitions are widespread and provide information beyond microbial abundance, their dynamics should not be reducible to changes in community or population size. Consistent with this expectation, activity and abundance exhibited distinct temporal patterns at both community and population levels, despite that their temporal stability were significantly and positively correlated (Fig. 3). At the community level, temporal stability in active-cell abundance was positively associated with redox activity stability (*R* = 0.59, *P* = 0.003; Fig. 3a), indicating a degree of coordination between community-level demographic and physiological dynamics after warming. However, interval-to-interval changes in active-cell abundance were overall weakly negatively correlated with changes in redox activity (*R* = −0.17, *P* = 0.038; Fig. 3b), but the strength and temporal pattern of this coupling varied across contexts. A multiple linear regression was conducted to examine whether this relationship is influenced by season and ecosystem type (adjusted *R*^2^ = 0.17). Analysis of variance (ANOVA) revealed a significant main effect of season (*F* = 11.96, *P* < 0.001), where the significant negative association was observed in both cold- and warm-season forest soils (slopes = −0.37 and −0.44, respectively) but was not detected in agricultural or grassland soils (95% CIs crossed zero).

**Fig. 3.**
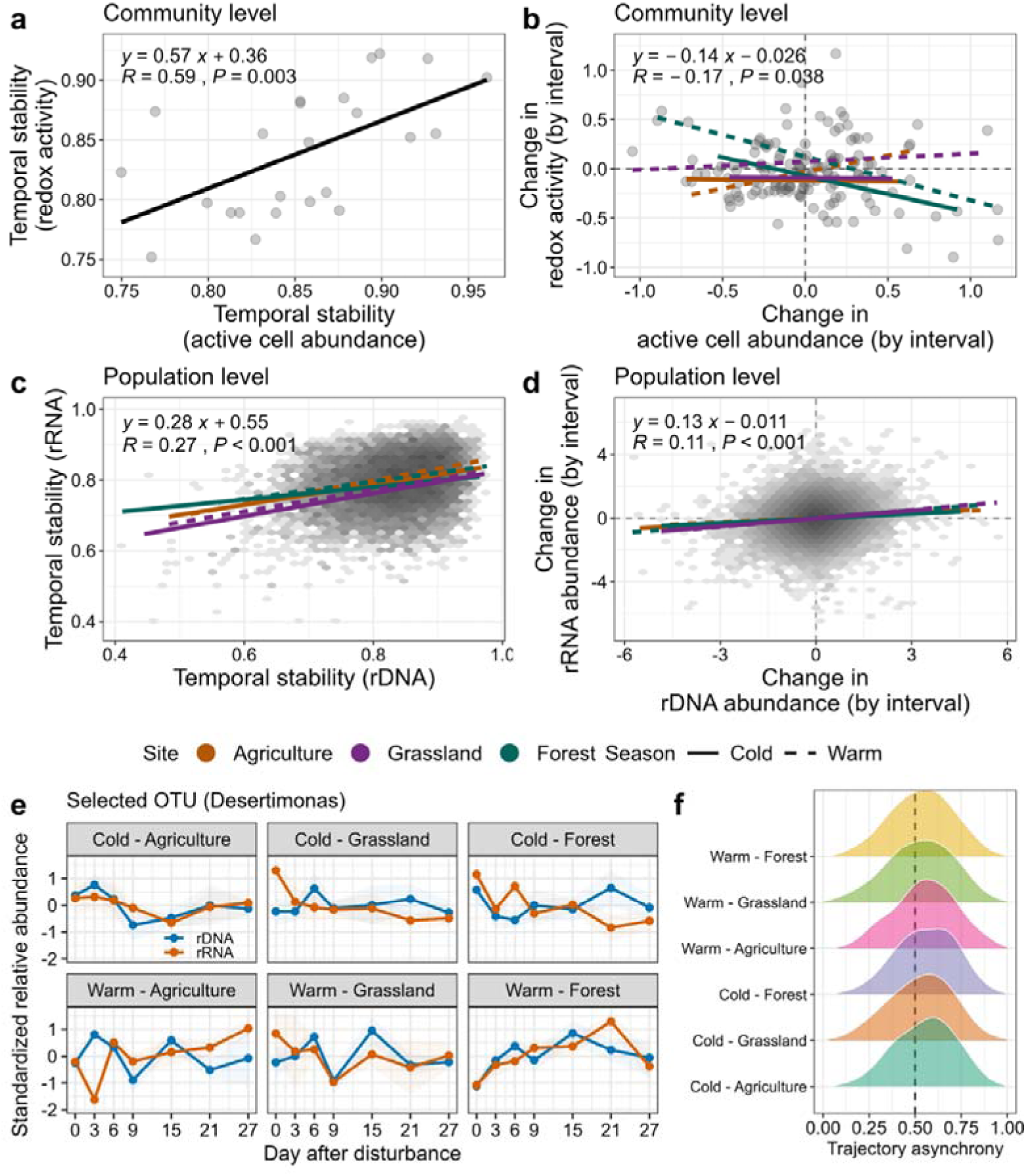
Asynchronous demographic and physiological dynamics at community and population levels. (**a**) Positive correlation between temporal stability of active cell abundance and redox activity, suggesting the coordination between community-level demographic and physiological dynamics after warming. (**b**) Community-level dynamic asynchrony as shown as the relationship between interval-to-interval changes in redox activity and active cell abundance. (**c-d**) Population-level relationships between demographic and physiological dynamics assessed by rDNA and rRNA abundance. Changes in abundance and activity indices at community and population levels are represented by the log2 fold change between two consecutive time points. Point density is indicated by grayscale shading, with darker shades representing higher OTU density. Regression lines are shown separately for agricultural soil, grassland, and forest from cold and warm seasons, with the statistical significance of these relationships and their differences were determined using a multiple linear model or linear mixed-effects models. The Pearson correlation coefficient (*R*) and significance of the overall association are shown within each panel. OTUs with an abundance of zero are not included in this analysis because the true log fold change cannot be precisely calculated. (**e**) An example of trajectory dynamics of a selected OTU (genus *Desertimonas*) across the six environmental conditions. (**f**) Trajectory asynchrony, represented by the normalized trajectory dissimilarity between rRNA and rDNA dynamics, showed broadly right-skewed distributions across all conditions, with the distributions extending predominantly towards higher asynchrony values relative to the mean value (vertical dashed lines).

At the population level, our testing revealed a positive association between the temporal stability of rDNA abundance and that of rRNA abundance (*R* = 0.27, *P* < 0.001; Fig. 3c). This relationship was further supported by a linear mixed-effects model incorporating random effects of taxonomic and microcosm-level variation (*P* < 0.001; marginal *R*^2^ = 0.086, conditional *R*^2^ = 0.39). Importantly, the relationship between rDNA and rRNA stability varied significantly among site types (*P* < 0.001) whereas seasonal differences did not significantly modify the relationship (*P* > 0.05). We found no evidence that this association varied across seasons, sites, or with their interaction (all interaction terms *P* > 0.05). In contrast, interval-level changes in rDNA abundance and rRNA abundance showed only a weak positive correlation (*R* = 0.11, *P* < 0.001; Fig. 3d). A linear mixed-effects model supported this association (*F* = 249.01, *P* < 0.001) and further suggested that this relationship varied among site types (*F* = 7.95, *P* < 0.001), whereas neither the interaction with season nor the three-way interaction between rDNA, season, and site type was significant (*P* = 0.20 and *P* = 0.17, respectively). Despite these significant associations, the model explained only a small proportion of the variation in rRNA changes (marginal *R*² = 0.013; conditional *R*² = 0.022), indicating that changes in rDNA abundance provided limited explanatory power for interval-level changes in rRNA abundance.

We further quantified the overall trajectory asynchronicity between response patterns of rRNA and rDNA and disentangled it from direction and magnitude of changes. The temporal trajectories of rRNA and rDNA abundance exhibited widespread asynchrony across the six environmental conditions (Fig. 3e-f), as illustrated by the selected OTU (*Desertimonas*). Consistently, trajectory asynchrony, represented by the normalized DTW distance between rRNA and rDNA trajectories, showed broadly right-skewed distributions across all six conditions (Fig. 3f), with the distributions extending predominantly towards higher asynchrony values relative to the mean value (0.5). This trajectory asynchrony differed significantly among site types and exhibited a significant season × site interaction (*P* = 0.002 and *P* < 0.001, respectively), whereas the overall seasonal effect was not significant (*P* = 0.17) after considering heteroscedasticity-robust standard errors (HC3). For instance, *post-hoc* comparisons revealed that trajectory asynchrony was significantly higher in cold forest than in warm forest (estimate = 0.076, Tukey-adjusted *P* = 0.004) and cold grassland (estimate = 0.086, Tukey-adjusted *P* = 0.0005).

Overall, these results indicate that temporal changes in microbial demographic and physiological responses were frequently asynchronous at the community and population levels, underscoring the importance of considering demographic and physiological processes as complementary dimensions of microbial community dynamics, as either dimension alone may provide an incomplete picture of microbial responses to environmental change.

### Initial physiological state structures early activity transitions

Warming subsequently induced extensive but context-dependent activity transitions across soil types and seasonal backgrounds (Fig. 4a-c and Supplementary Text 2). Forest communities showed a pronounced excess of activation over deactivation, particularly in cold-season soils, whereas agricultural communities showed the opposite tendency; grassland communities displayed stronger seasonal shifts in the balance between activation and deactivation (Fig. 4a). These transitions were also taxonomically structured, with several bacterial classes showing contrasting activation and deactivation patterns among soil types (Supplementary Results Text 2). Importantly, forest communities contained a particularly large contribution of populations transitioning toward higher activity, including initially low ribosomal activity populations, whereas agricultural communities showed greater representation of deactivation trajectories (Fig. 4c).

**Fig. 4.**
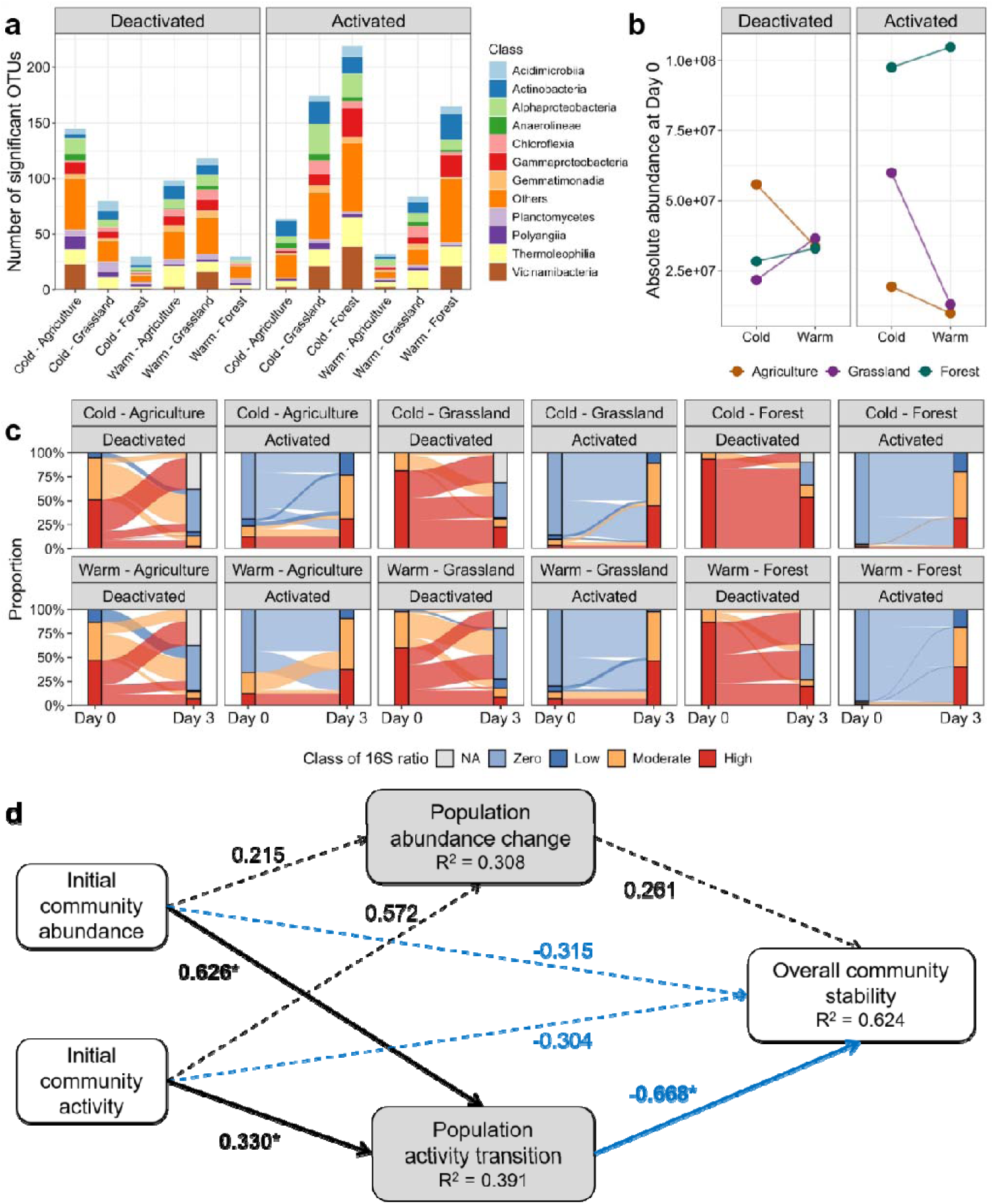
Deciphering population activity transitions for predicting community stability under disturbance. (**a**) Numbers of OTUs showing significant changes in physiological activity (16S rRNA:16S rDNA, also known as 16S ratio) between Day 0 and Day 3, classified as deactivated or activated according to the direction of change. Bars are partitioned by bacterial taxonomic class. (**b**) Cell abundance at Day 0 of OTUs showing significant deactivation or activation, summarized by site type and season. (**c**) Transitions in 16S-ratio states from Day 0 to Day 3 for OTUs showing statistically significant deactivation or activation. OTUs were classified as NA (not detected in rDNA dataset), Zero, Low (16S ratio < 0.25), Moderate (0.25 ≤ 16S ratio < 1), or High (16S ratio ≥ 1) according to their 16S ratio, and the width of each flow represents the proportion of OTUs transitioning between states. (**d**) PLS-SEM showing standardized path coefficients for the effects of initial community states (Day 0) on population activity and abundance transitions (Day 0-3, composited with count, proportion, and initial abundance of OTUs) and overall community stability integrated with abundance, activity, and composition (Day 0-27). Solid arrows indicate significant (\**P* < 0.05) and dashed indicate non-significant paths (*P* > 0.05) with the standardized path coefficients. Blue arrows denote negative effects, and black arrows denote positive effects. Full model specifications are provided in Supplementary Table 2.

The early activity transitions of individual OTUs were strongly structured by their initial physiological states. Initial 16S ratio at Day 0 was negatively associated with subsequent changes in 16S ratio over the first 3 days (slope = −0.21, *R* = −0.34, *P* < 0.001, Supplementary Fig. 5). OTUs with higher initial ribosomal activity therefore tended to decrease their activity, whereas those with lower initial ribosomal activity tended to increase it (Supplementary Fig. 6), indicating a consistent convergence toward intermediate physiological states. This phenomenon was also observed in redox activity at the community level. In contrast, initial absolute abundance showed marginal explanatory power for subsequent activity changes (slope = 0.036, *R* = 0.084, *P* < 0.001, Supplementary Fig. 5). Thus, early activity transitions were more closely structured by the initial physiological state of individual populations than by their initial abundance. Together, these patterns indicate that early physiological transitions were shaped jointly by the initial activity state of populations and by site- and season-specific environmental context.

Abundance responses nevertheless involved substantial population turnover and varied markedly among soil types and seasonal backgrounds, with distinct sets of bacterial populations increasing or decreasing across environmental contexts (Supplementary Figs. 7 and 8 and Supplementary Results Text 2). In contrast to activity transitions, early changes in population abundance were only weakly structured by initial states. Initial ribosomal activity and initial abundance showed little explanatory value for subsequent changes in absolute abundance over the first 3 days (slope = 0.045 and −0.037, *R* = −0.087 and −0.1, respectively; *P* < 0.001 and 0.001; Supplementary Figs. 5). Thus, unlike the pronounced dependence of early activity transitions on initial physiological state, early abundance changes were comparatively weakly predictable from the initial demographic or physiological state of populations.

### Early activity transitions and resistance predict community and population stability

The PLS-SEM explained a substantial portion of the variance in transitional dynamics and overall community stability (Fig. 4c). The model explained 62.4% of the variation in overall community stability (*R*^2^ = 0.624) that contains multiple dimensional responses such as community cell abundance, redox activity, and composition. Initial community abundance and redox activity were negatively correlated with overall stability, but not significant, with path coefficients (β) of −0.315 and −0.303, respectively (*P* > 0.05). Notably, population activity transitions exhibited a stronger negative association with stability (β = −0.668, *P* < 0.05), highlighting the importance of dynamic activity responses in shaping subsequent community stability. The model also explained 39.1% of the variation in population activity transitions (*R*^2^ = 0.391), suggesting that early community properties may help explain how microbial populations adjust their activity following warming. In comparison, population abundance change was explained 30.8% of the variation (*R*^2^ = 0.308) and was not significantly associated with the community stability (β = 0.261, *P* > 0.05, Fig. 4c). These results highlight a potential cross-scale link between microbial population dynamics and community stability, suggesting that early shifts in population-level physiological activity may provide insights into the stability of the broader microbial community.

We next examined how initial community states and early resistance (Days 0–3) were associated with the subsequent stability of community composition, active-cell abundance, and community activity during Days 3–27 (Supplementary Fig. 9 and Table 2). Initial community abundance and activity were both negatively associated with early resistance in activity (β = −0.910 and −0.356, *P* < 0.05), indicating that the magnitude of the initial state alone did not necessarily translate into a more resistant community. Moreover, initial community activity was negatively associated with early resistance in community active-cell abundance (β = −0.581, *P* < 0.05), indicating the fundamental role of microbial activity in community changes. Notably, early activity resistance was associated with the subsequent stability of community composition (β = 0.767, *P* < 0.05), whereas abundance-related metrics showed no comparable associations (Supplementary Fig. 9). None of the examined predictors showed significant direct associations with the subsequent stability of community active-cell abundance or redox activity independently (Supplementary Fig. 9). These findings suggest that early physiological responses may provide information about the persistence of community composition that is not captured by abundance-based metrics alone, highlighting the potential importance of activity dynamics in linking immediate disturbance responses to longer-term community stability.

A similar pattern emerged at the population level, where the later stability of population abundance and ribosomal activity was more strongly associated with initial activity states or early activity resistance than with abundance-related metrics (Supplementary Fig. 10 and Table 2). Together, these results suggest that microbial physiological activity and its early responses to disturbance provide indicative information about subsequent community dynamics that is not captured by abundance alone.

## 4 Discussion

Climate change can have strong effects on microbial growth and alter community composition, with consequences for ecosystem processes and functioning^37^. Few studies have examined how microbial activity responds to disturbance while also examining how it relates to subsequent community outcomes like stability. This study shows that early physiological activity transitions provide predictive information about later microbial community stability following persistent warming. Across soils differing in ecosystem types and seasonal history, activity transitions were distinct from abundance changes, structured by initial physiological state, and associated with subsequent population and community stability.

Microbial abundance and physiological activity are previously considered to be interconnected and even positively correlated because population growth ultimately requires the conversion of physiological activity into biomass. Advances in our understanding of microbial physiological states, particularly dormancy and the capacity of microorganisms to maintain substantial biomass under low-activity conditions^38,39^, have highlighted that standing microbial biomass does not necessarily reflect the physiologically active fraction of the community. This recognition has motivated increasing efforts to distinguish microbial abundance from physiological activity at both population and community levels. For instance, Metze *et al.* (2024)^16^ have suggested that increased bacterial community growth under warming was associated primarily with an increase in the number of growing taxa, rather than higher growth rates across populations. Conversely, Söllinger *et al.* (2024)^40^ found that warming consistently reduced community-level 16S ratios across seasons and warming durations, indicating an overall adjustment in ribosome investment that occurred without corresponding shifts in community composition. These studies suggest a potential decoupling of the positive mathematical correlation between microbial abundance and physiological activity. Our results extend this emerging view by resolving the temporal dynamics of these two dimensions following disturbance at both population and community levels in natural soil communities. Importantly, we found that the temporal stability of demographic and physiological metrics tended to be positively associated after warming, suggesting the coordinated responses to environmental disturbance across these two dimensions. However, their short-term dynamics were much less tightly coupled over the same period, implying that the maintenance of population or community abundance does not necessarily require a consistent level of physiological activity, and that changes in physiological state can occur without proportional changes in population size. Such a temporal separation is consistent with the ability of microorganisms to modulate metabolic investment, ribosome allocation, and cellular maintenance without immediately altering population size, as observed during transitions between high-activity and low-activity states^38,41–43^. In this sense, microbial activity transitions may provide a physiological buffer against environmental perturbation^44^, allowing populations to adjust their level of resource acquisition and cellular investment while delaying the demographic consequences of that adjustment. Consequently, incorporating activity measurements provides an additional dimension for interpreting microbial responses that cannot be inferred from abundance alone.

The ecological implications of demographic and physiological asynchrony may differ between population and community levels, reflecting distinct processes ranging from physiological regulation within individual populations to compositional turnover and compensatory dynamics at the community level. At the population level, individual microbial taxa may adjust their physiological activity in response to environmental fluctuations before detectable changes in population abundance occur. For example, a recent study showed that changes in rRNA abundance in an aquatic microbial community precede changes in rDNA as seasonal and environmental oscillations dynamically change^45^. It might be similar in soil communities, especially given the general sensitivity of microbes to temperature^9^. At the community level, however, demographic and physiological asynchrony may additionally emerge from shifts in the relative contributions of different taxa, including the activation or suppression of specific populations and compensatory changes among taxa with overlapping functional capacities. These scale-dependent processes suggest that microbial responses to environmental disturbances are unlikely to be uniform across taxa or ecosystems. Rather, the magnitude and direction of demographic and physiological responses may depend on taxon-specific ecological strategies and pre-existing environmental conditions, including resource availability, environmental history, and the composition and functional organization of the resident microbial community.

Indeed, our results showed that the apparent convergence in community- and population-level physiological activity dynamics did not imply that the ecological mechanisms or outcomes at the community level are consistent. Despite initial convergence in community-level activity among soils, physiological activity subsequently fluctuated significantly over time, with the amplitude influenced by both season and ecosystem type. Similarly, the populations undergoing activation or deactivation differed strongly among ecosystem types and season histories, both in number and in their initial abundance and activity states. Forest communities, for example, contained a larger pool of populations with initially low activity that could be activated following warming, including transitions from undetected or low-activity states to higher activity states. These low-activity populations may therefore represent a physiological “seed bank”^19^ that is not readily apparent from conventional abundance-based measurements, but that can become functionally important when environmental conditions change. This finding suggests that the response capacity of a community is determined not only by which populations are present, but also by the distribution of their physiological states before disturbance. Therefore, microbial activity transitions provide a mechanistic link between community history and ecosystem response: environmental history shapes the reservoir of physiological states available for activation or deactivation, while disturbance redistributes populations along this activity spectrum. The ecological consequences of disturbance may thus depend on whether microbial activity changes, which populations undergo these activity transitions, the initial physiological states of those populations, and their abundances.

Early physiological transitions, estimated by changes in 16S rRNA-to-rRNA gene ratio (also known as rRNA:rDNA ratio), contained information about subsequent stability that abundance changes did not. Population activity transitions were strongly associated with overall community stability, and initial activity states and early activity resistance predicted later stability at both population and community levels. These transitions reveal how populations redistribute across physiological states, linking environmental history and the initial state structure of a community to its subsequent trajectory. This interpretation is consistent with cross-site evidence that climate and soil properties shape microbiome vulnerability to extreme events^46,47^ and with functional regimes that link metabolic activity to community and ecosystem responses^48^. By resolving the direction, magnitude, and prevalence of population-level activity transitions, our framework may therefore identify changes in community stability before they are fully manifested in abundance or composition.

While our controlled microcosm approach successfully isolated the fundamental microbial dynamics under persistent warming, natural environments introduce additional ecological complexities, such as fluctuating climates and plant-microbe interactions, that can ultimately modulate these responses. Furthermore, although our experimental design captured distinct contrasting land-use and seasonal contexts, expanding this framework to broader climatic regions will help establish the global generality of these patterns. It is also important to contextualize our activity metrics: while the rRNA:rDNA ratio and RSG-based measurements act as specific indices for ribosomal potential and reductase activity rather than absolute metabolic flux, these data suggest that they effectively capture the overall physiological shifts driving community trajectories. Future experiments that directly manipulate microbial activity-state distributions will be vital to unraveling the precise causal mechanisms underpinning these dynamics and integrating these physiological indicators into broader ecosystem models.

To conclude, our findings highlight that microbial physiological responses are a necessary dimension for detecting community dynamics and support a predictive framework in which early physiological activity transitions link environmental history to later community stability under persistent warming disturbance. This work presents an advance in understanding how soil microbiomes will respond and their capacity to stabilize with the changing climate.

## Acknowledgements

This work was funded by the European Union (ERC, MicroRescue, 101087042). Views and opinions expressed are, however, those of the authors only and do not necessarily reflect those of the European Union or the European Research Council. Neither the European Union nor the granting authority can be held responsible for them. We thank Estelle Couradeau, Juliana Almario, J. Gregory Caporaso, and William R. Shoemaker for constructive comments on this work; Ludivine Guigard and Wei Zheng for soil sampling; Emma Chavan for molecular biology experiments; and Jonathan Gervaix and the AME (Microbial Activities in the Environment) platform for soil respiration experimental support. We acknowledge the IN2P3 Computing Centre (CC-IN2P3) for providing the computing and storage resources used in this work.

## Author contributions

A.S. conceived the research and secured funding for the project. A.S. and X.L. designed the experiment and wrote the original draft. X.L. and A.F.B. conducted the microcosm experiment with significant contributions from M.G. and M.B. on the laboratory work. X.L. performed data analysis and visualization. All authors contributed to editing and finalizing the draft.

