## Supplementary_Materials for "Microbial activity transitions underlie community stability following persistent warming"

This manuscript includes Supplementary Methods, Supplementary Text 1-2, Fig. 1-10, and Table 1-2.

### Supplementary Methods

#### Soil microbial abundance and redox activity

**Cell separation and extraction.** Soil cells were separated with a modified density gradient separation protocol based on ours and others’ previously published protocols^1,2^. In brief, 6 g of soil were mixed with 25 mL of extraction buffer (50 mM tetrasodium pyrophosphate with 0.5% Tween80, TSP-T) and blended at high speed for 1 min. After pelleting large soil particles (centrifuged at 1000 × g for 10 min), the soil suspension from 1 g of soil was calculated accordingly and slowly transferred on top of 4 mL of 1.0 g/mL Nycodenz solution (Serumwerk Bernburg AG, Germany) and then centrifuged for 40 min at 12000 × g under 25 °C. Then, 2 mL of the upper phase of Nycodenz containing microbial cells into a sterile 2 mL tube. The second extraction was immediately performed for the pelted soil particles mentioned above by mixing them with 25 mL of TSP-T buffer, following the steps in the first extraction until obtaining soil microbial cells. We then combine the cells from two extractions by centrifuging them at 12,000 × g for 10 min at the origin temperature of soil samples and resuspending those cells in 1 mL of TSP buffer. Intact cells in the soil are thus considered to have been extracted.

**Cell counting and abundance.** To assess the number of total and active cells and its redox activity, the extracted fresh cells were diluted and stained with SYBR^TM^ Green I (Invitrogen, CA, USA), which stains DNA-containing cells^2^, and Redox Sensor Green (RSG, BacLight RedoxSensor green vitality kit, Invitrogen), which binds to microbial reductases^3,4^, respectively. In details, 10 μL of diluted cell suspension was mixed with 1 μM of SYBR or 1 μM of RSG with an incubation at room temperature in the dark for 15 minutes. Later, 6 μL of the stained cell mixture was loaded to a loading slide with the same volume of 100% glycerol and centrifuged at 650 × g for 10 min in a specific-slide centrifuge (QUANTOMTM^TM^ Centrifuge, Q10002) to immobilize and uniformly distribute the cells on the slide. The automated microscopy was performed using the QUANTOM microbial cell counter (Q100001, Logos Biosystems 2019), which captures cells visible with an excitation of 470/30 nm and an emission of 530/50 nm (Logos Biosystems Inc., South Korea). This cell counter captured ten discrete images from each loading slides. A developed pipeline (process_quantom: <https://github.com/DatInsights-com/process_quantom>) was used to detect individual cells in images and then capture variables of interest, such as cell counting, florescence intensity, etc^1^. The cell abundance was presented as cell count per gram of dried soil.

**Single-cell redox activity and community activity.** In all images obtained for each sample, the RSG fluorescence intensity of all pixels recognized as an individual cell was summed as the total RSG fluorescence intensity, representing the redox activity of the cell. We then consider three measures of community redox activity. First, the median redox activity is the median total RSG fluorescence intensity of all active cells (RSG-positive) in each individual community. Second, the cumulative redox activity of a community is calculated as the summed total RSG fluorescence intensity of all active cells (RSG-positive), calculated as the fluorescence intensity per gram of dry soil, representing the redox capacity of the whole community. Third, the mean redox activity is the normalized cumulative activity by the total cell count (SYBGreen-positve).

#### Microbiota profiling

**DNA and RNA co-extraction.** Soil samples were added to 2 mL RNase/DNase-free microtubes containing 0.7 mm garnet and 0.1 mm glass beads. 0.5 ml of lysis solution (5% CTAB, 120 mM phosphate buffer at pH 8, and 1.6 M NaCl), and 0.5 ml of phenol:chloroform:isoamyl alcohol (25:24:1) were added to the tube. Cells were lysed via bead beating for 60 seconds, followed by centrifugation at 12,000 g for 10 minutes at 4 °C. The resulting aqueous phase was recovered and further purified by adding an equal volume of chloroform:isoamyl alcohol (24:1), vortexing, and centrifuging at 16,000 g for 5 minutes at 4 °C. Total nucleic acids were precipitated from the final aqueous layer by adding two volumes of a PEG precipitation solution (30% PEG6000 and 1.6 M NaCl) and incubating on ice for 2 hours. After centrifugation at 16,000 g for 30 minutes at 4 °C, the pellet was washed with 1 ml of ice-cold 70% ethanol. The purified pellet was air-dried to avoid ethanol residue, resuspended in 50 µl of DEPC-treated water, and quantified using a Qubit fluorometer. The extract was then aliquoted into two fractions: one for DNA storage at -80 °C and another for RNA processing, including DNase treatment and subsequent cDNA synthesis.

**DNase treatment and cDNA synthesis.** To obtain pure RNA, DNA was removed using the TURBO DNA-free Kit (Thermo Fisher Scientific). Extracts were diluted to approximately 40 ng/µL of DNA. A 50 µL reaction mixture—comprising 25 µL of diluted extract, 5 µL of 10X Turbo DNase Buffer, 3 µL of Turbo enzyme, and 21 µL of nuclease-free water—was incubated at 37°C for 30 minutes. The reaction was terminated by adding 2 µL of DNase Inactivation Reagent, followed by a 5-minute incubation at room temperature with periodic mixing. The purified RNA was transferred to a fresh tube and the absence of genomic DNA contamination was confirmed via gel electrophoresis.

Reverse transcription was performed using the GoScript™ Reverse Transcription System (Promega, France). Briefly, 4 µl of purified RNA was primed with Random Primer and incubated at 70°C for 5 minutes followed by 5 minutes on ice. A reaction mix containing GoScript™ 5X reaction buffer, MgCl_2_, PCR nucleotide mix, and GoScript™ Reverse Transcriptase was added to a final volume of 20 µL. The synthesis conditions consisted of annealing at 25°C for 5 minutes, extension at 42°C for 1 hour, and enzyme inactivation at 70°C for 15 minutes. The resulting cDNA was used immediately for 16S rRNA V4 gene amplification to confirm the possibility of amplification and then stored at -80°C for downstream use.

**Illumina sequencing of 16S rRNA and 16S rRNA gene.** The V3-V4 region of the 16S rRNA gene was amplified from extracted DNA and purified cDNA using the universal bacterial and archaeal primers 341F (5′- CCTACGGGAGGCAGCAG-3′) and 806R (5′-GGACTACHVGGGTWTCTAAT-3′). PCR amplification was performed in 25-μL reactions containing Q5^®^ High-Fidelity master mix (New England Biolabs), 0.2 μM of each primer, and average 2-4 ng of template. Thermal cycling consisted of an initial denaturation at 94°C for 3 min, followed by 25 cycles of 94°C for 30 s, 53°C for 40 s, and 72°C for 60 s, followed by a final extension at 72°C for 5 min. Each sample was amplified in duplicate independent PCR reactions, and replicate reactions were pooled prior to purification. Amplicons were purified using Sera-Mag™ Carboxylate-Modified Magnetic Beads (Cytiva), quantified using Quant-iT™ PicoGreen™ dsDNA Analysis Kits (Invitrogen, CA, USA), and subjected to a second 10-cycle PCR to incorporate dual-index barcodes and Illumina sequencing adapters. Indexed libraries were purified, quantified, normalized, and pooled at equimolar concentrations. The final library was sequenced on an in-house Illumina NextSeq 1000 using 2 × 300 bp paired-end sequencing chemistry. Negative PCR controls were processed alongside samples throughout library preparation to monitor potential contamination.

**Bioinformatics.** Paired-end FASTQ files were imported into QIIME 2 using the PairedEndFastqManifestPhred33V2 format. Primer sequences targeting the V3–V4 region were removed using Cutadapt, with untrimmed reads discarded. Taxonomic classification of representative sequences was performed using a Naive Bayes classifier trained on the SILVA 138.2 reference database. DNA- and cDNA-derived sequences were classified independently, after which sequences and features assigned to mitochondria, chloroplasts, or eukaryote were removed. Potential contaminants were identified independently in the DNA and cDNA datasets using the prevalence-based *decontam* approach, with negative controls used to estimate contaminant probabilities. Features with a contamination probability >0.5 were removed from subsequent analyses. Following contaminant removal, the DNA and cDNA feature tables and representative sequences were merged to enable consistent feature definition across the two molecular fractions. The combined dataset was clustered de novo at 99% sequence identity using VSEARCH, generating operational taxonomic units (OTUs) for downstream analyses. OTU clustering was preferred over ASV inference to reduce the influence of fine-scale sequence variation on population matching between DNA- and cDNA-derived communities^5^. This provided a consistent operational definition of microbial populations for subsequent comparisons of abundance and activity. Taxonomy was reassigned to the resulting OTU representative sequences using the SILVA 138.2 classifier, and the OTU abundance and taxonomy tables were exported for downstream statistical analyses in R. Repeated random rarefaction was performed at a depth of 14,714 sequences per sample with 100 independent iterations using the “BRCore” package^6^ (v. 2.0.7). The mean abundance of each OTU in each sample was calculated across the 100 rarefaction iterations.

### Supplementary Results

#### Text 1: Detailed responses of microbial abundance, activity, and community composition

*Resistance and temporal stability of total cell abundance*

Resistance of total cell abundance showed significant effects of season and ecosystem type at both Day 3 and Day 27, with significant season × site type interactions in both cases (Day 3: season, *P* = 0.0063; site type, *P* < 0.001; interaction, *P* < 0.001; adjusted *R*² = 0.73; Day 27: season, *P* = 0.0021; site type *P* = 0.024; interaction, *P* = 0.0017; adjusted *R*² = 0.61, Supplementary Fig. 1). Pairwise comparisons revealed pronounced ecosystem-specific differences in cold-season soils. At Day 3, resistance was higher in agriculture than forest (*P* < 0.0001) and in forest than grassland (*P* = 0.0001), whereas agriculture and grassland did not differ significantly (*P* = 0.13). A similar pattern persisted at Day 27, with higher resistance in agriculture than forest (*P* = 0.0002) and in forest than grassland (*P* = 0.011), while agriculture and grassland remained statistically indistinguishable (*P* = 0.20). In contrast, none of the pairwise comparisons among ecosystem types was significant in warm-season soils at either Day 3 or Day 27 (*P* > 0.58). Thus, resistance of total cell abundance exhibited strong ecosystem-specific differentiation under cold-season soils, whereas these differences were largely absent under warm-season soils. For the temporal stability of total cell abundance, neither season nor ecosystem type had a significant effect, and their interaction was also nonsignificant (season, *P* = 0.19; site type, *P* = 0.073; season × site type, *P* = 0.33; adjusted *R*² = 0.19). Although mean stability was highest in agriculture (0.876), followed by grassland (0.842) and forest (0.812), none of the pairwise differences remained significant after Tukey adjustment (all *P* > 0.06). Thus, unlike resistance, temporal stability of total cell abundance showed no clear differentiation among seasons or ecosystem types.

*Resistance and temporal stability of active cell abundance*

Resistance of active cell abundance exhibited distinct temporal patterns across seasonal history and soil types (Fig. 2a). At Day 3, resistance was significantly affected by season (*F* = 7.41, *P* = 0.014), site type (*F* = 6.88, *P* = 0.0060), and their interaction (*F* = 7.76, *P* = 0.0037), with the model explaining 67.1% of the variation (adjusted *R*² = 0.58). For communities collected from cold season, resistance was higher in agriculture than forest soils (estimate = 0.24, *P* = 0.004) and higher in grassland than forest soils (estimate = 0.34, *P* = 0.0002), whereas agriculture and grassland soils did not differ (*P* = 0.33). In contrast, no significant differences among soil types were detected in warm-season conditions (all *P* ≥ 0.87), indicating a strong soil-type dependence of resistance under cold conditions that was largely absent under warm conditions. By Day 27, season (*F* = 5.26, *P* = 0.034) and site type (*F* = 11.08, *P* = 0.0007) remained significant predictors of resistance, whereas the season × site type interaction was no longer significant (*F* = 0.065, *P* = 0.94). Under cold conditions, agriculture and grassland soils continued to exhibit higher resistance than forest soils (*P* = 0.025 and 0.014, respectively), while no significant pairwise differences were detected under warm conditions (all *P* ≥ 0.05). In contrast to resistance, temporal stability of active cell abundance was comparatively consistent across seasons and site types (Fig. 2a). Neither season (*F* = 1.78, *P* = 0.20), site type (*F* = 2.82, *P* = 0.086), nor their interaction (*F* = 1.68, *P* = 0.21) significantly affected stability (adjusted *R*² = 0.20). Pairwise comparisons similarly revealed no significant differences among soil types in the cold season (*P* > 0.93).

*Resistance and temporal stability of community redox activity*

Resistance of community median redox activity was strongly structured by site type at Day 3 (*F* = 32.95, *P* < 0.0001), whereas the effect of season was not significant (*F* = 2.42, *P* = 0.14); the season × site type interaction was marginal (*F* = 5.68, *P* = 0.012, adjusted *R*² = 0.77, Fig. 2b). Under both cold and warm conditions, forest soils exhibited lower resistance than grassland soils (estimate = −0.12, *P* = 0.0068; estimate = −0.20, *P* < 0.0001), while the agriculture–forest contrast was not significant (*P* > 0.05). By Day 27, both site type (*F* = 6.19, *P* = 0.009) and its interaction with season (*F* = 8.30, *P* = 0.002) significantly affected resistance, while the main effect of season remained non-significant (*P* = 0.40, adjusted *R*² = 0.52). Notably, no differences among site types remained in the cold season (*P* > 0.78), whereas in the warm season agricultural soils exhibited higher resistance than both forest and grassland soils (estimate = 0.27 and 0.18, respectively; both *P* < 0.01). Median redox stability showed an even stronger and more consistent dependence on environmental context. Stability was significantly affected by Season (*F* = 23.40, *P* = 0.00013), Site type (*F* = 23.41, *P* < 0.0001), and their interaction (*F* = 3.38, *P* = 0.056), with the model explaining 81.0% of the variation (adjusted *R*² = 0.76). Under cold conditions, agriculture exhibited higher stability than forest (estimate = 0.051, *P* = 0.035), while forest was less stable than Grassland (estimate = −0.083, *P* = 0.0008); Under warm conditions, agriculture again showed higher stability than forest (estimate = 0.10, *P* < 0.0001) and grassland (estimate = 0.031, *P* = 0.25), while forest remained less stable than grassland (estimate = −0.073, *P* = 0.0026).

#### Text 2: Site- and season-specific transitions of activity and abundance

Warming elicited widespread changes in microbial activity, but the magnitude and direction of these responses varied substantially across ecosystem types and seasonal backgrounds (Fig. 4a-c). At the physiological level, deactivation and activation involved markedly different numbers of OTUs across sites and seasons (Fig. 4a). The number of deactivated OTUs was highest in agriculture (149 OTUs), intermediate in grassland (80), and lowest in forest (25) among soils collected during the cold season, whereas the corresponding numbers were 98, 119 and 30 OTUs, respectively, among soils collected during the warm season. In contrast, activated OTUs increased from 59 in agriculture to 173 in grassland and 219 in forest among cold-season communities, compared with 31, 82 and 165 OTUs, respectively, among warm-season communities. Thus, forest communities showed a particularly strong excess of activation over deactivation, whereas agriculture showed the opposite tendency, with grassland displaying a stronger seasonal shift.

The taxonomic composition of activity transitions revealed distinct soil- and season-specific patterns across bacterial classes. For instance, Actinobacteria, Vicinamibacteria and Gammaproteobacteria showed a particularly strong contrast, with a large number of activated OTUs in both cold and warm forest soils but comparatively few deactivated OTUs, whereas agricultural soils contained substantially more deactivated than activated Vicinamibacteria and Gammaproteobacteria populations. These differences were also reflected in the initial abundance of responsive OTUs (Fig. 4b). Deactivated OTUs were generally more abundant in cold agricultural soils (6.3 × 10^7^ cells), whereas their abundance was more evenly distributed among site types in the warm season. In contrast, activated OTUs were particularly abundant in forest soils in both cold and warm seasons (9.8 × 10^7^ and 1.05 × 10^8^ cells), with their abundance remaining high in warm forest soils while declining sharply in agricultural and grassland soils.

Early abundance responses involved substantial numbers of bacterial populations and differed markedly among soil types and seasonal backgrounds (Supplementary Fig. 8). Among cold-season communities, the number of significantly decreased OTUs ranged from 65 in forest soils to 446 in grassland soils, while significantly increased OTUs ranged from 84 in agricultural soils to 623 in forest soils. Cold-season forest communities therefore showed a pronounced predominance of OTUs with increased abundance, whereas cold-season grassland communities contained the largest number of significantly decreased OTUs. These patterns indicate that warming-associated changes in population abundance were highly heterogeneous among soil types, even within the same seasonal background. These abundance shifts involved multiple bacterial classes, with several taxonomic groups contributing prominently to both decreased and increased populations. The major contributors included Gammaproteobacteria, Gemmatimonadia, Alphaproteobacteria, Actinobacteria and Acidimicrobiia, together accounting for a substantial fraction of the significantly changing OTUs. The taxonomic composition of increased and decreased OTUs therefore differed across environmental contexts, indicating that disturbance-associated population turnover was distributed across diverse bacterial lineages rather than being restricted to a single taxonomic group.

### Supplementary Figures

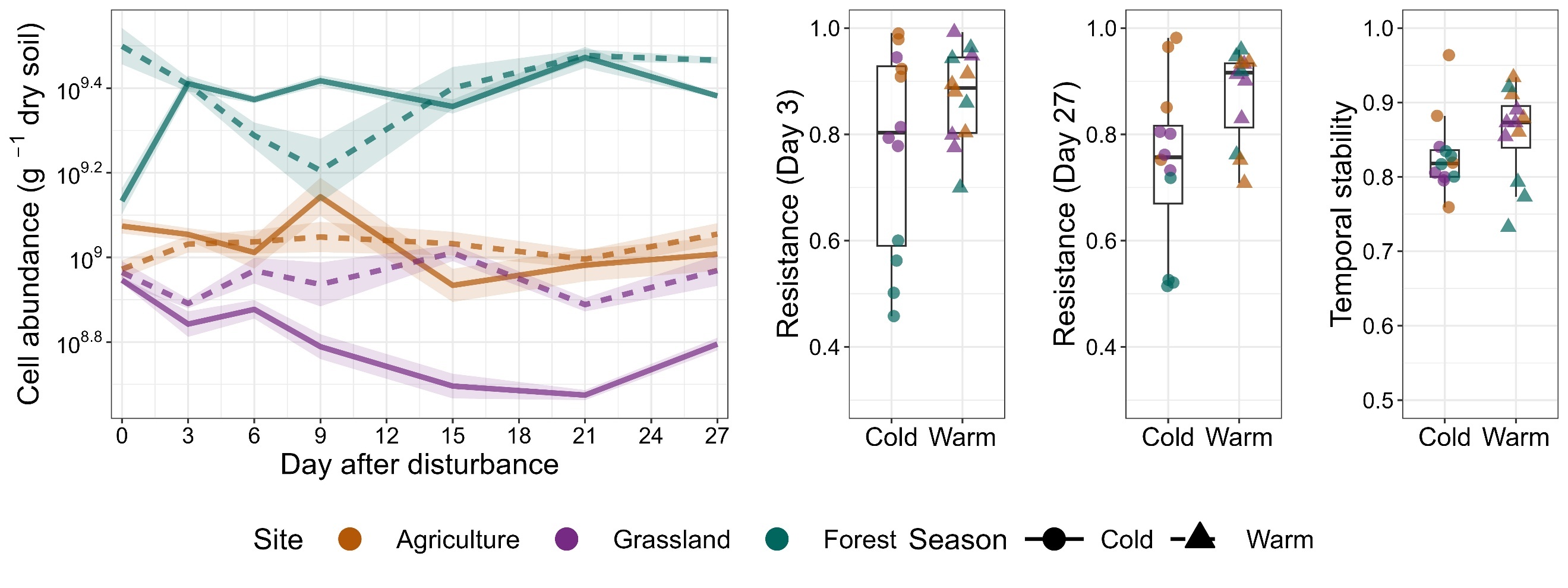

S. Fig. 1 Dynamic, resistance, and temporal stability of total cell abundance in soils. The shading around the line represents the standard error (n=4 microcosms). Cold and warm denote the two seasonal conditions of soil origin, whereas agriculture, grassland, and forest represent ecosystem types.

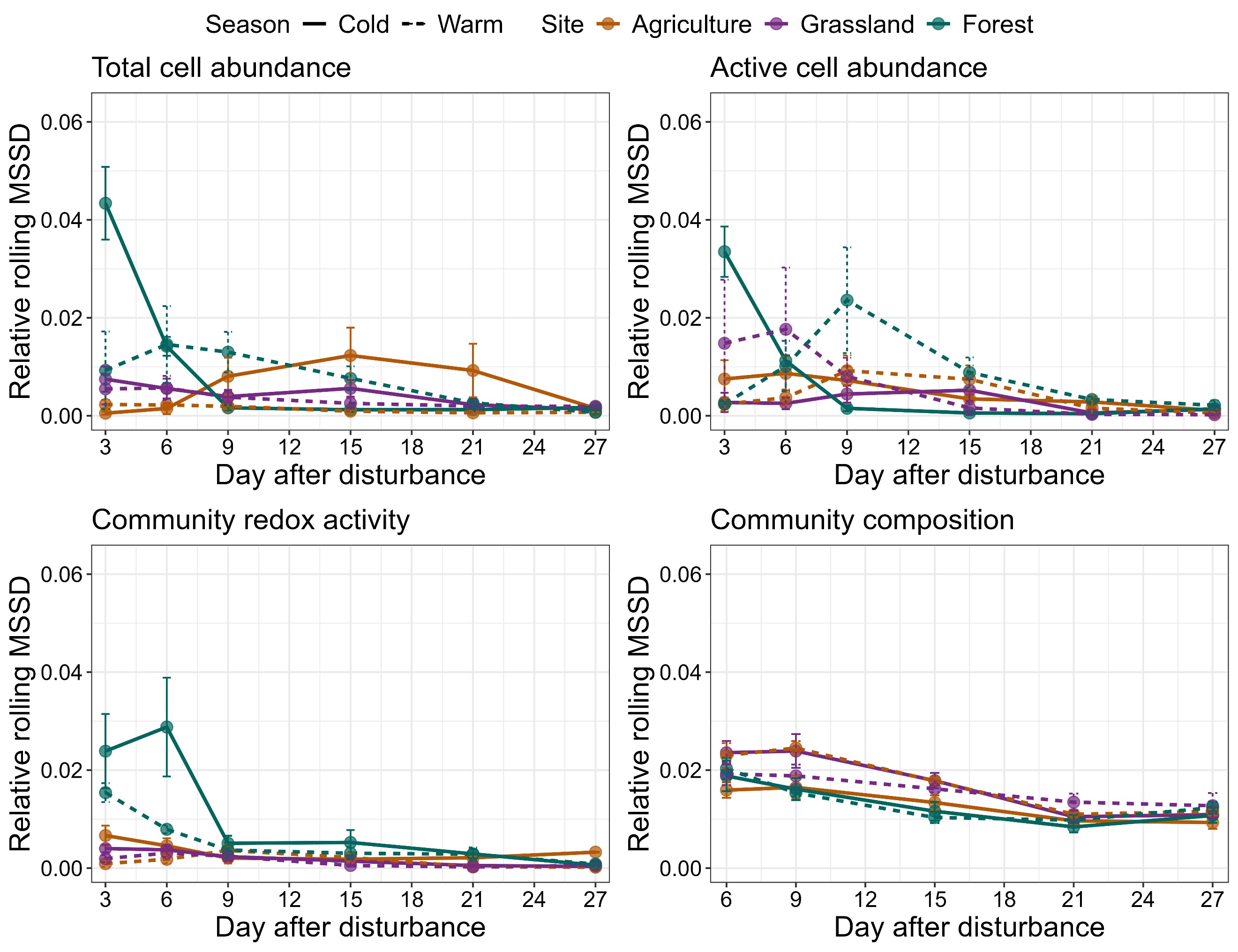

S. Fig. 2 Short-term temporal variability and compositional turnover following disturbance across seasons and site types. Relative rolling mean squared successive difference (MSSD) for total cell abundance, active-cell abundance, community redox activity, and community composition (rolling succession rate) over 27 days after disturbance. MSSD is a dimensionless index of temporal instability (window = 2 observations). Lines are group means ± SE; colors denote site type and line type denotes season. Instability peaked early (Days 3–9) and declined thereafter, with forest soils showing the highest initial variability.

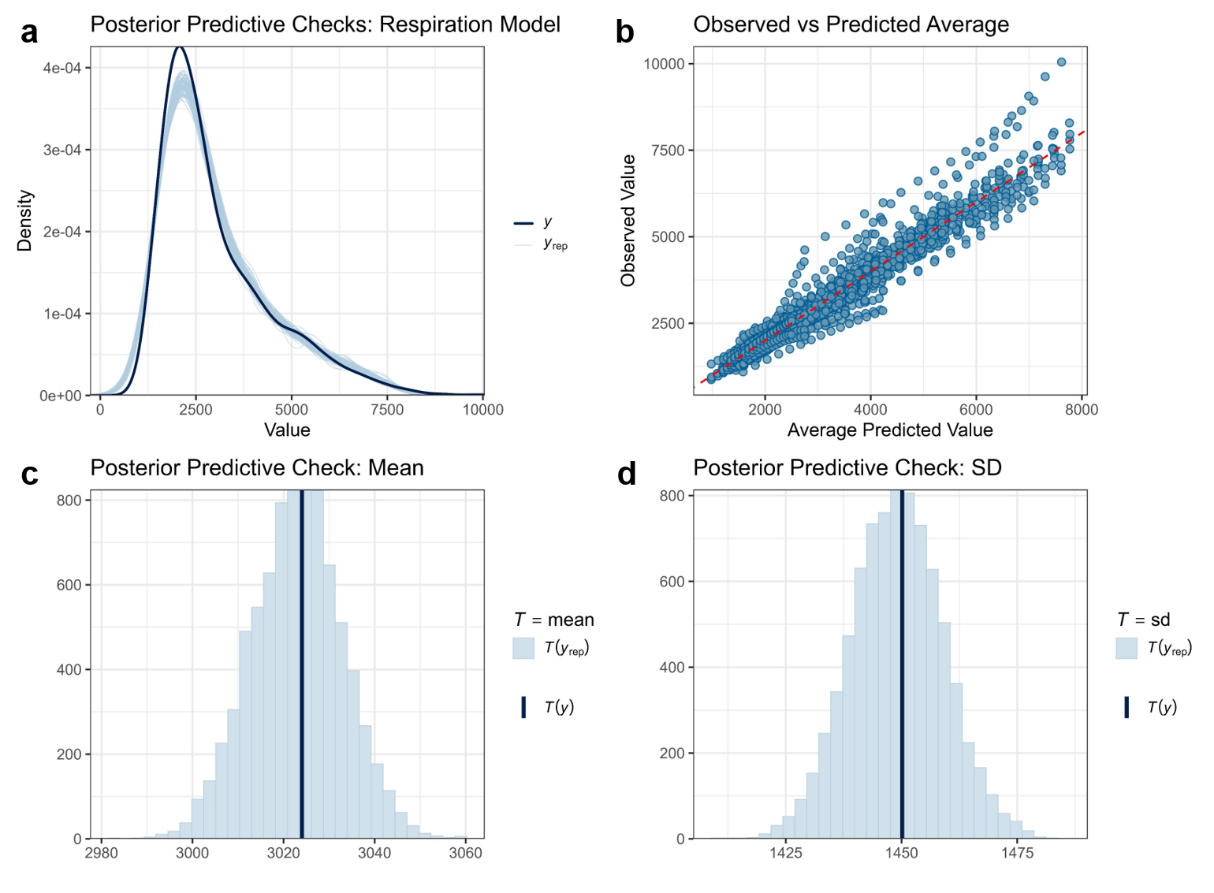

S. Fig. 3 Posterior predictive checks for the Bayesian soil respiration model. (a) Comparison of the observed respiration distribution with distributions simulated from the posterior predictive distribution. (b) Observed versus posterior-predicted average values; the red dashed line indicates the 1:1 relationship. (c) Comparison of the observed mean with the distribution of means calculated from posterior predictive replicates. (d) Comparison of the observed standard deviation (SD) with the distribution of SDs calculated from posterior predictive replicates. These diagnostics evaluate the model's ability to reproduce the observed distribution, mean, and variability of soil respiration.

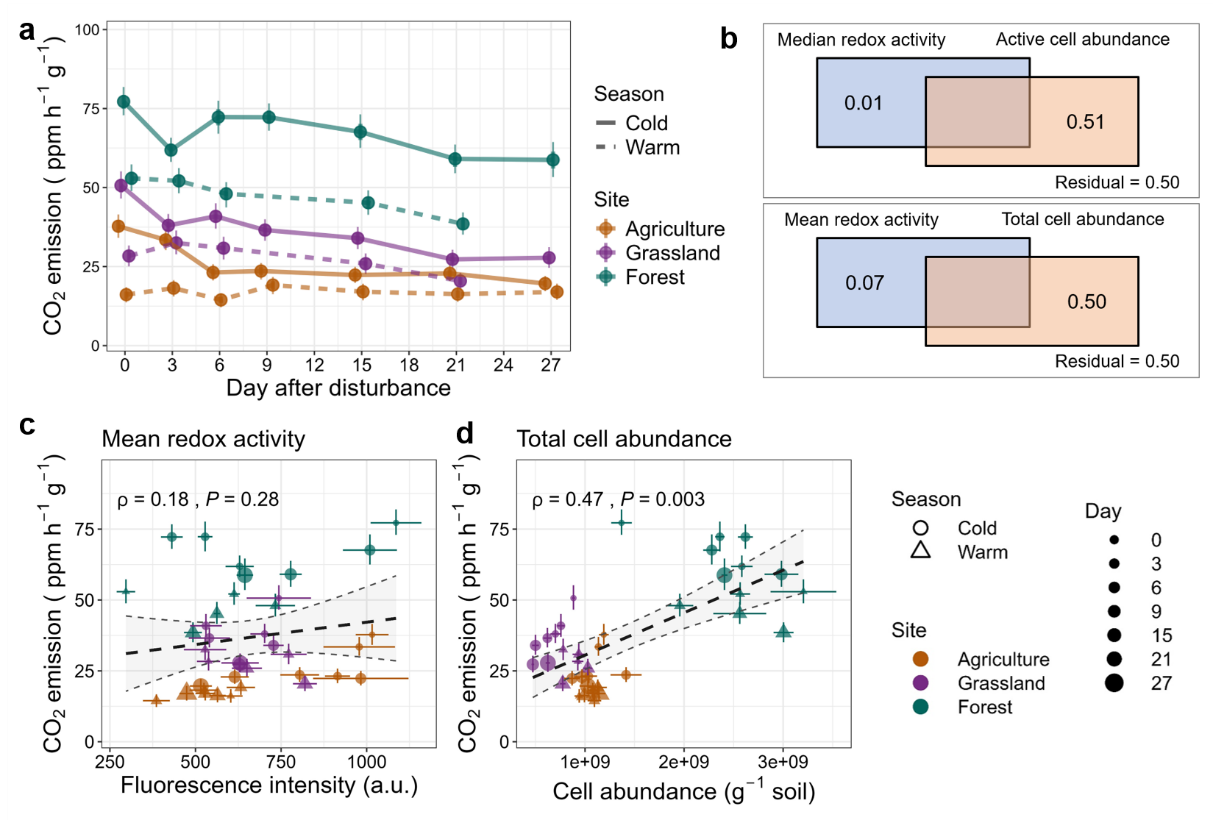

S. Fig. 4 (a) Soil CO₂ emissions capacity. Points represent posterior median estimates of CO₂ emission, with thick and thin intervals indicating the 66% and 95% highest-density credible intervals (HDIs), respectively. Lines show median CO₂ emission across sampling days, with line type indicating season and color indicating site type. (b) Variance partitioning analysis (VPA) quantifying the contributions of microbial physiological activity and community size to initial soil respiration. Mean redox activity and total cell abundance, or median redox activity and active-cell abundance, were used as activity and size metrics, respectively. The absence of an overlap indicates no detectable shared explanatory contribution between community activity and community size. (c-d) Relationships between multiple dimensions of microbial community properties and soil CO₂ emission following disturbance. (c) Mean cellular redox activity normalized by total cell abundance, (d) total cell abundance. Spearman’s correlation was shown as regression lines and corresponding 95% confidence intervals. Points represent median posterior estimates of CO₂ emission and mean estimates of cumulative microbial activity. Vertical error bars indicate 95% credible intervals of CO₂ emission derived from the posterior distribution, whereas horizontal error bars represent ±1 standard error of microbial community properties (n=4).

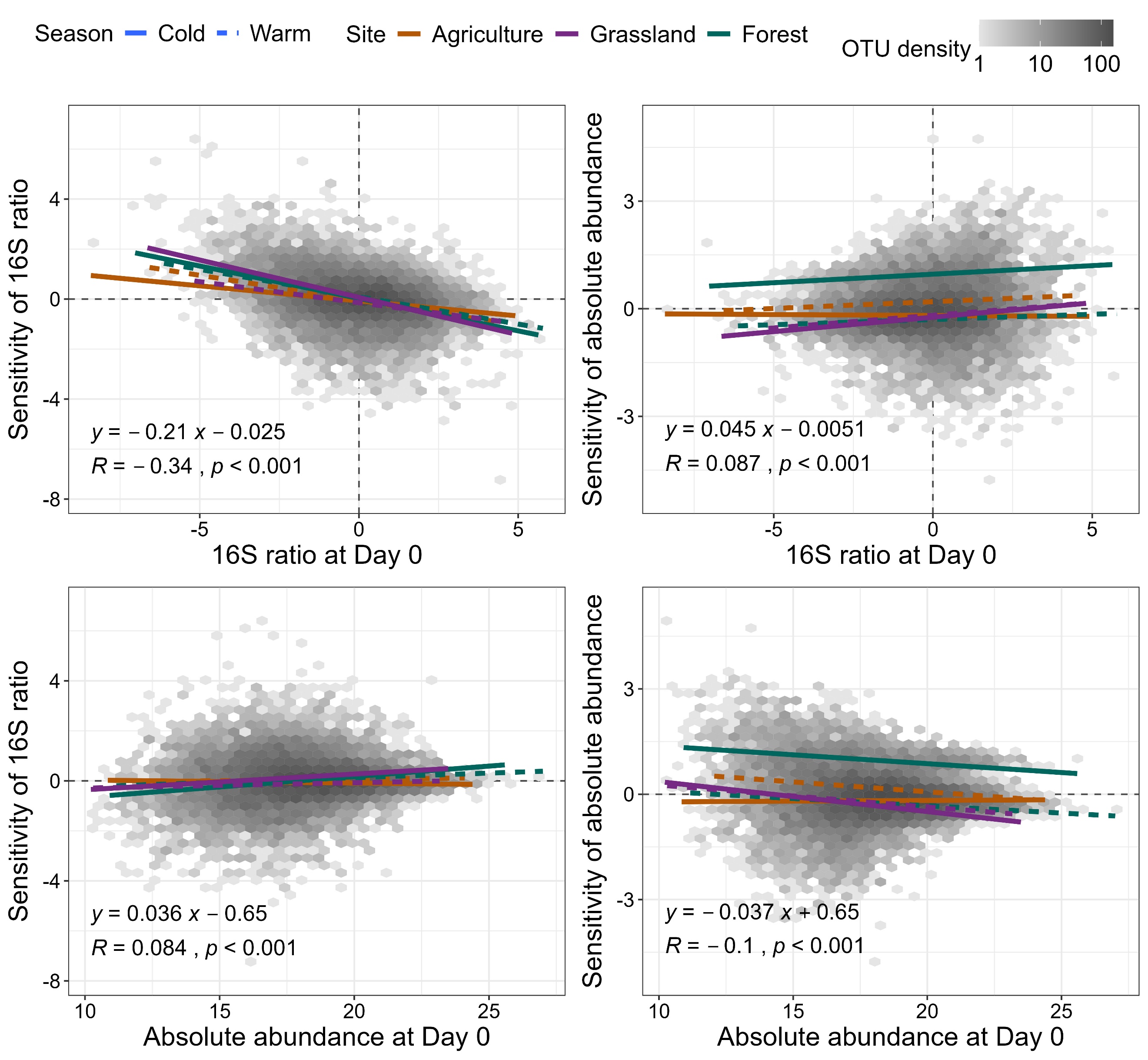

S. Fig. 5 (a) Correlation between initial OTU states and changes in population activity and absolute abundance at Day 3. Sensitivity was calculated as the log2 fold change in 16S ratio and cell abundance from Day 0 to Day 3 with a positive value representing an increase. The x-axis represents the log2-transformed initial 16S ratio (upper panels) and absolute abundance (lower panels) of OTUs at Day 0.

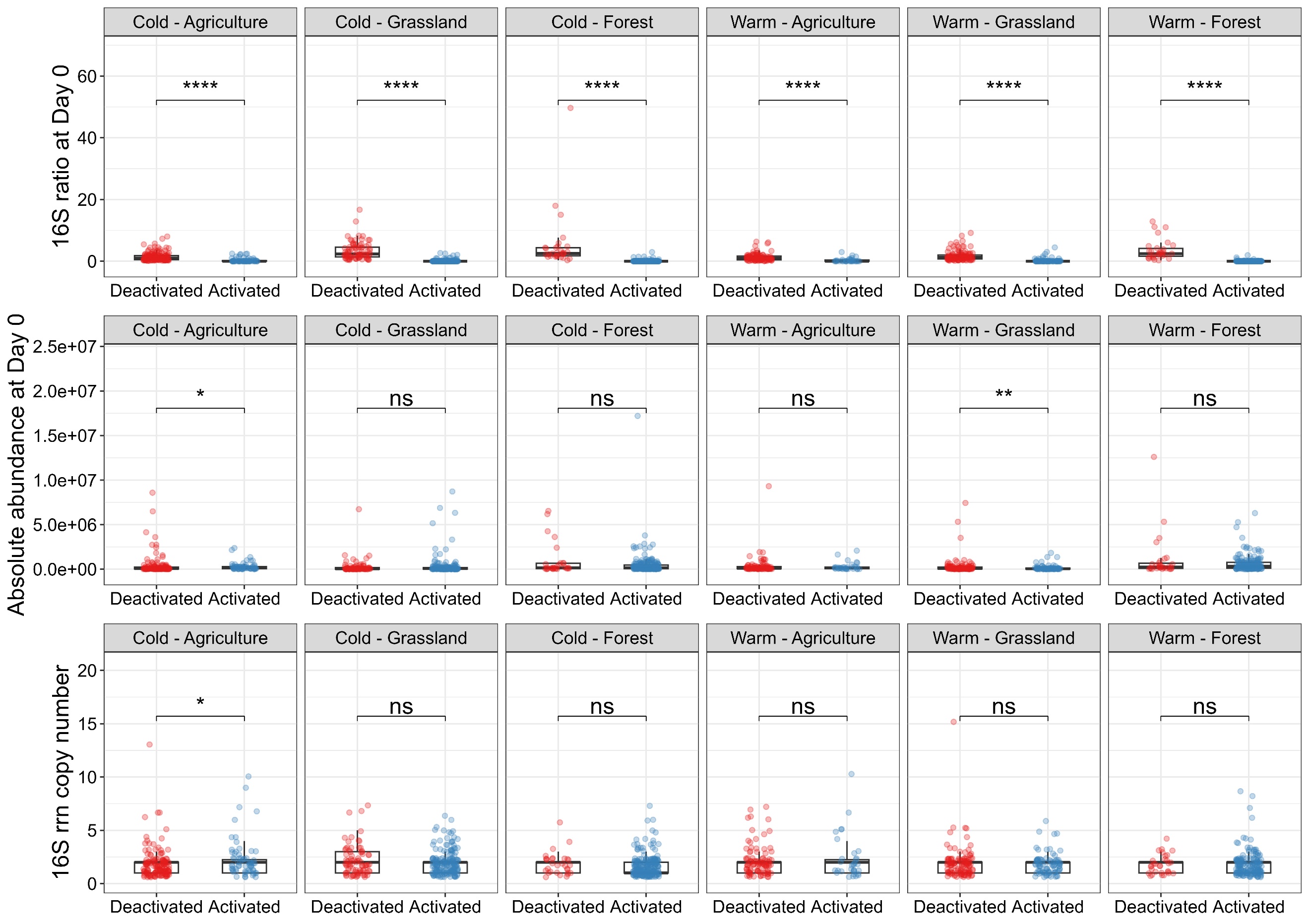

S. Fig. 6 Comparison of two categories of OTUs that were significantly changed in their 16S ratios between Day 0 and Day 3. The two categories were deactivated and activated according to the direction of change. Differences between activated and deactivated populations were assessed using the Wilcoxon rank-sum test (* *P* < 0.05, ** *P* < 0.01, *** *P* < 0.001, **** *P* < 0.0001).

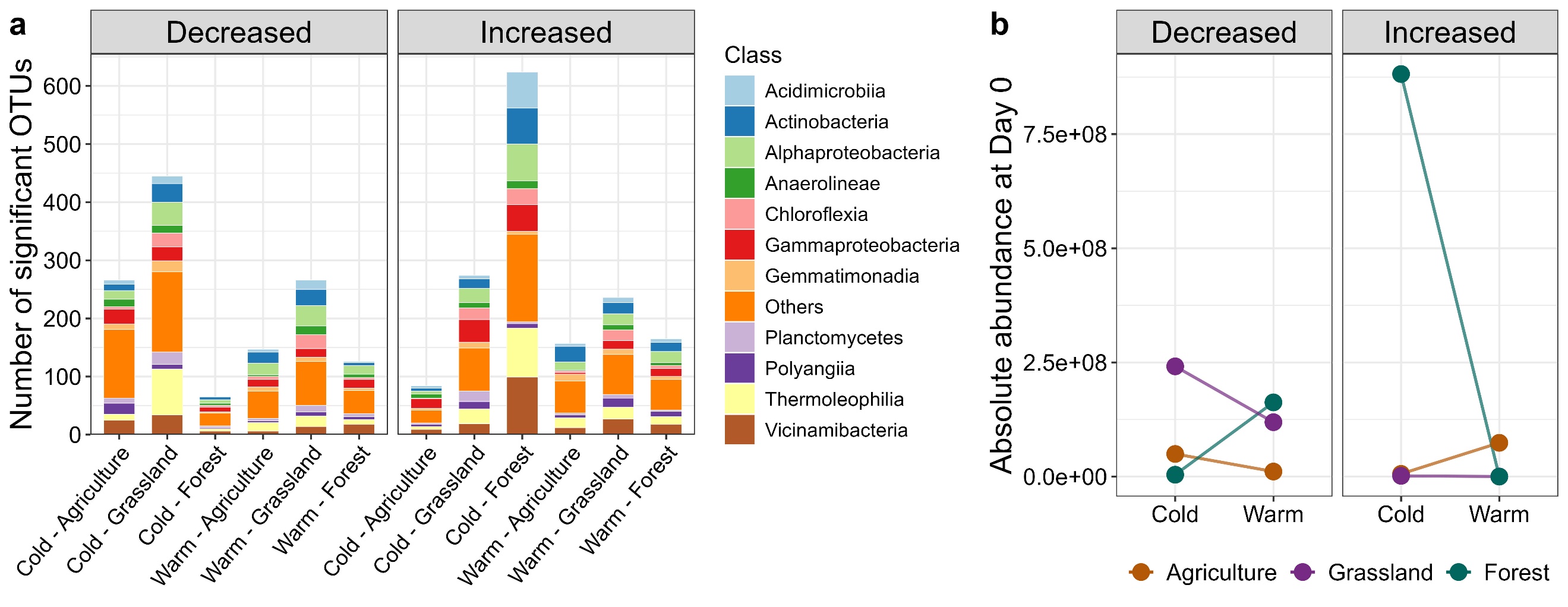

S. Fig. 7 Abundance responses to persistent warming. (a) Numbers of OTUs showing significant decreases or increases in OTU absolute abundance between Day 0 and Day 3, with bars partitioned by bacterial taxonomic class. (b) Initial absolute abundance of OTUs (Day 0) showing significant decreases or increases in absolute abundance, summarized by ecosystem type and season. Cold and warm denote the two seasonal conditions of soil origin, whereas agriculture, grassland, and forest represent ecosystem types.

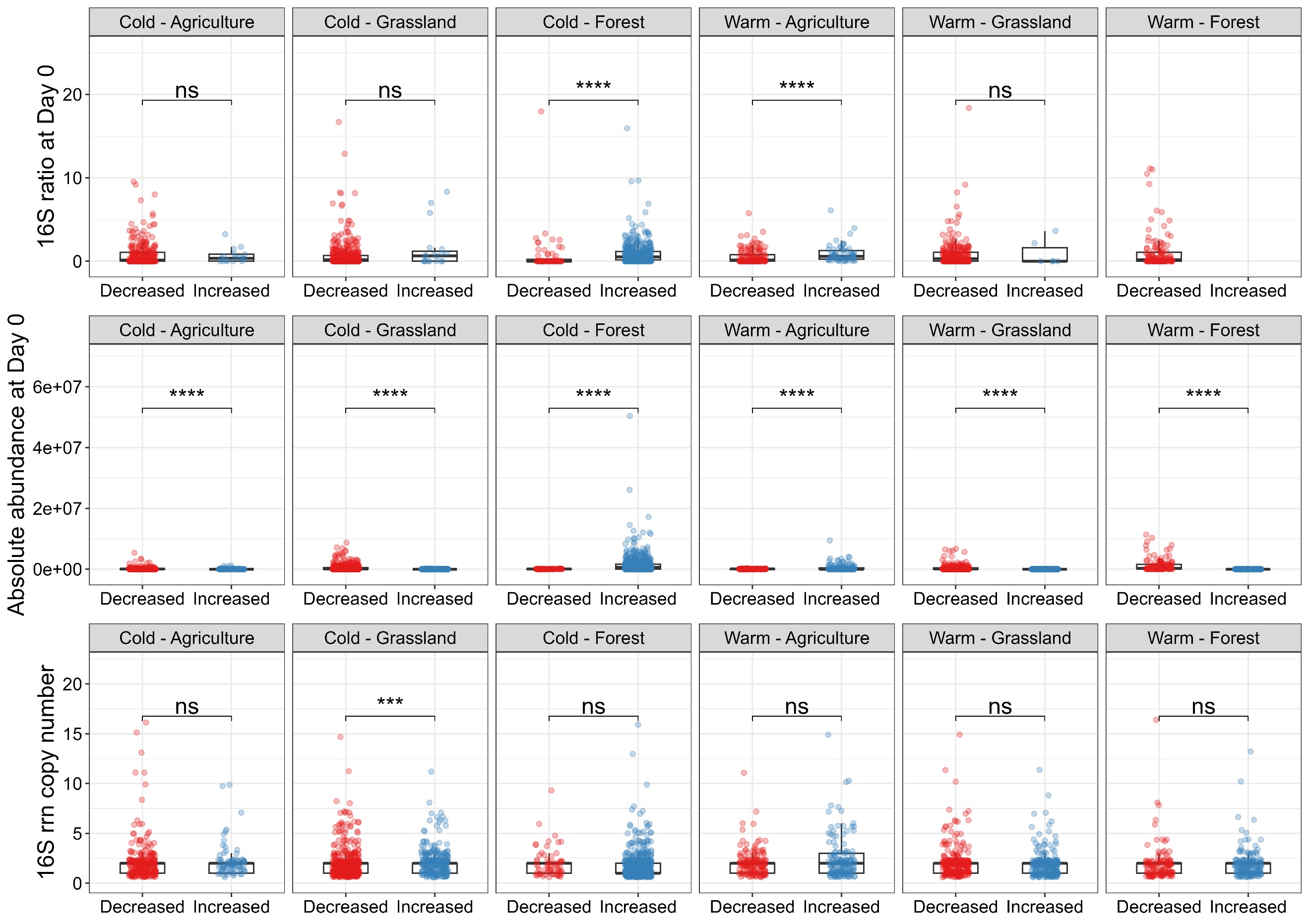

S. Fig. 8 Comparison of two categories of OTUs that were significantly changed in their absolute abundance between Day 0 and Day 3. Two categories were decreased and increased according to the direction of change. Differences between activated and deactivated populations were assessed using the Wilcoxon rank-sum test (* *P* < 0.05, ** *P* < 0.01, *** *P* < 0.001, **** *P* < 0.0001).

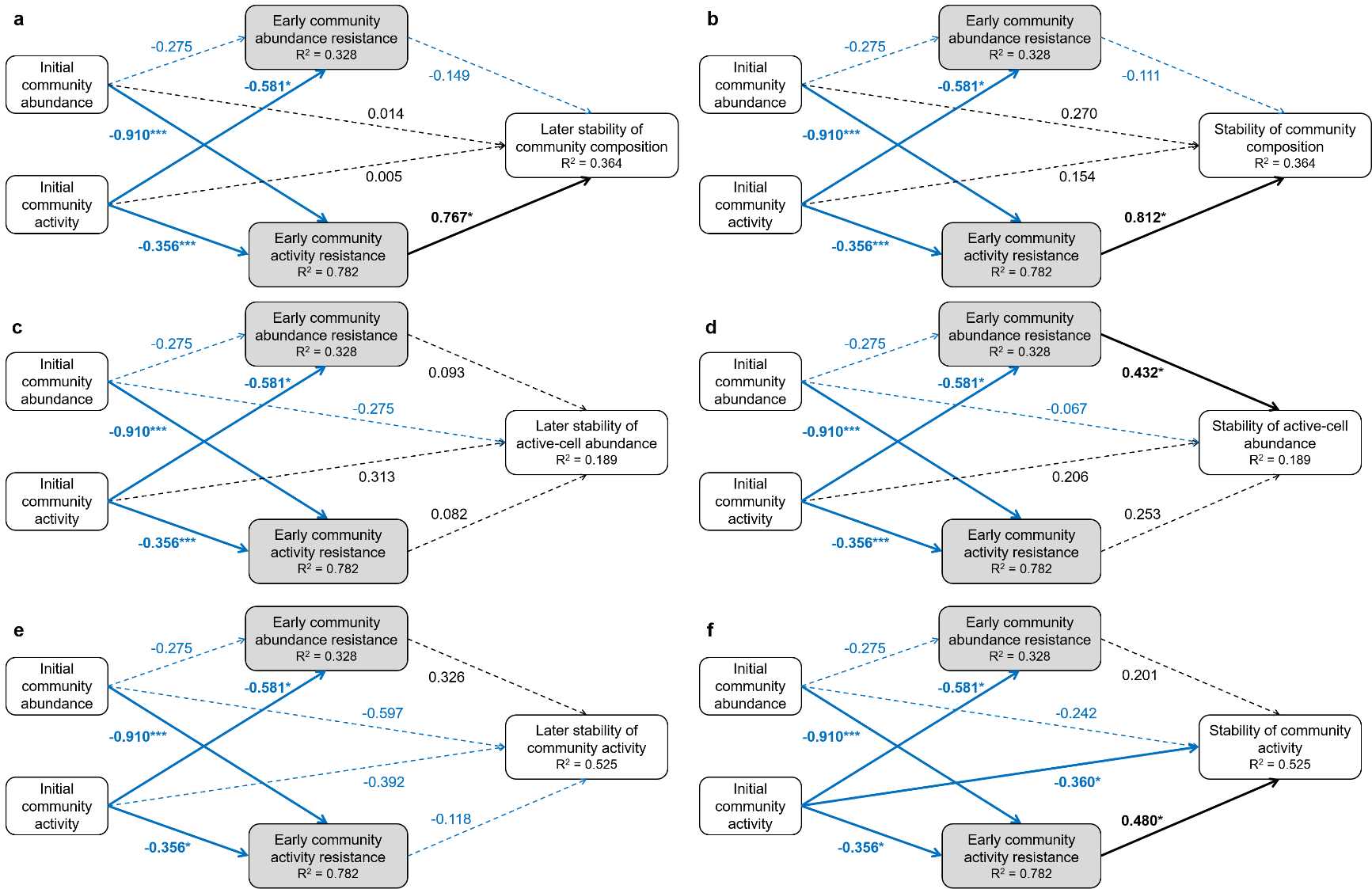

S. Fig. 9 PLS-SEM showing standardized path coefficients for the effects of initial community states (Day 0) on early community resistance (Day 0-3), later (Day 3-27) and overall (Day 0-27) stabilities of community composition (a-b), active-cell abundance (c-d), and redox activity (e-f). Note that each panel has the same input parameters (initial active-cell abundance, redox activity, early resistance in cell abundance and redox activity) but differ in their objective variables. Solid arrows indicate significant (**P* < 0.05, *P* < 0.01, ****P* < 0.001) and dashed arrows indicate non-significant (*P* > 0.05) paths with the standardized path coefficients. Blue arrows denote negative effects, and black arrows denote positive effects. Full model specifications are provided in Supplementary Table 2.

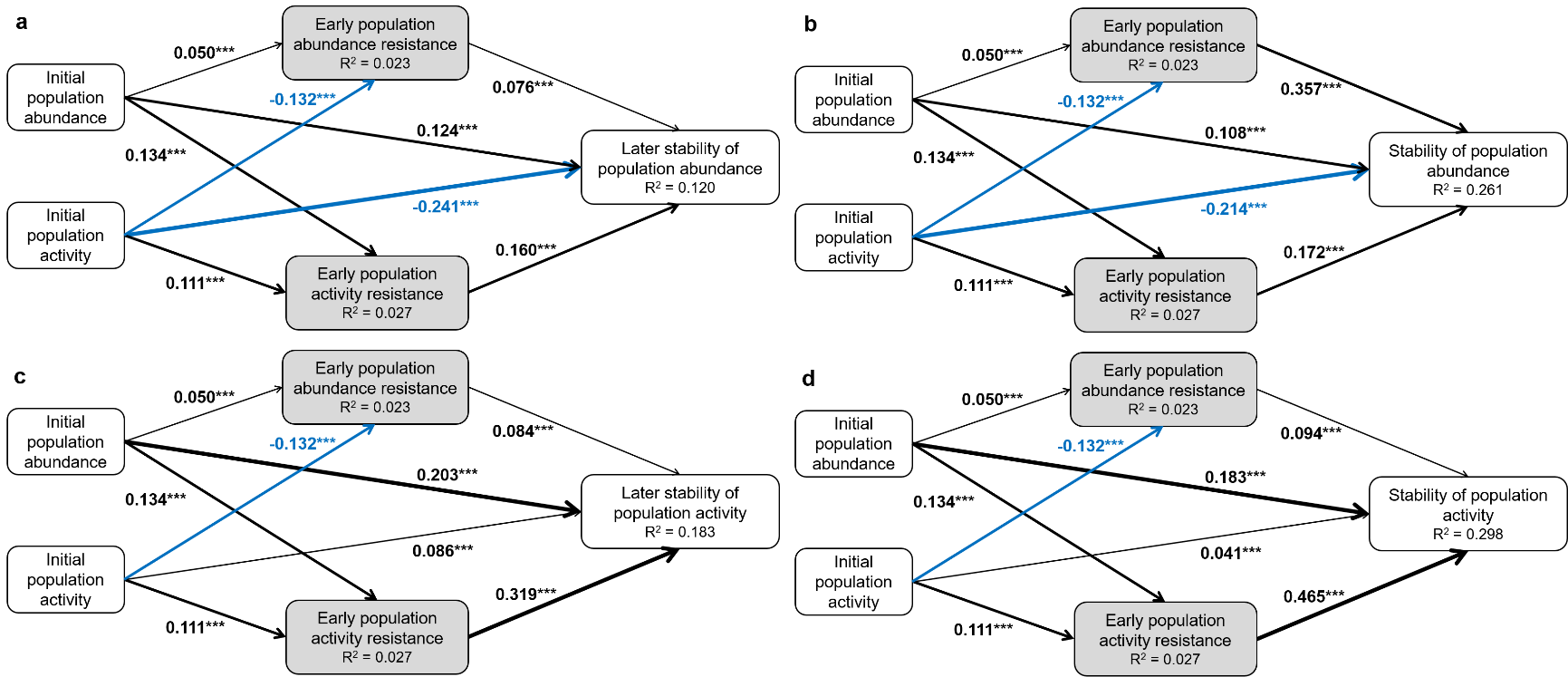

S. Fig. 10 PLS-SEM showing standardized path coefficients for the effects of initial population states (Day 0) on population resistance (Day 0-3) and later (Day 3-27) and overall (Day 0-27) population stabilities of with (a-b) absolute cell abundance and (c-d) ribosomal activity (16S ratio). Arrows are indicating significant (**P* < 0.05, *P* < 0.01, ****P* < 0.001) path with the standardized path coefficients. Blue arrows denote negative effects and black arrows denote positive effects. Full model specifications are provided in Supplementary Table 2.
